# Accurate and efficient prediction of protein conformations with ProtMonomer

**DOI:** 10.64898/2026.08.28.747824

**Authors:** Yunda Si, Suqi Zhang, Luonan Chen

## Abstract

Deep learning-based protein structure prediction methods that leverage evolutionary information from multiple sequence alignments (MSAs), exemplified by AlphaFold2, have achieved remarkable accuracy. However, existing methods still struggle to predict challenging proteins, particularly those with novel folds or limited evolutionary information, and to recover alternative conformational states. Here we show that structure prediction models trained under different MSA-depth distributions corresponding to different levels of evolutionary information exhibit complementary generalization behaviors, and that a model trained on a mixture of these distributions can combine their complementary generalization strengths. Building on this insight, we developed ProtMonomer, a deep learning framework trained on MSA-depth distributions representing a broad range of evolutionary information levels to improve structure prediction. Across benchmarks comprising CASP15 targets, non-redundant experimentally determined structures, orphan proteins, and short peptides, ProtMonomer performed comparably to or better than leading methods, including AlphaFold2 and AlphaFold3, with particularly strong performance on challenging targets. For fold-switching proteins, ProtMonomer also recovered alternative conformational states more accurately than AlphaFold2 and AlphaFold3 across diverse homologous sequence sampling strategies. In addition to improving predictive accuracy, ProtMonomer substantially reduced inference cost through an efficient architecture, enabling high-throughput applications. Together, these findings provide insights into the generalization of evolution-informed structure prediction models and support ProtMonomer as an accurate and efficient framework for protein structure prediction.

## Introduction

Accurate modeling of three-dimensional protein structures is essential for understanding protein function and advancing rational protein design^1,2^. Evolutionary information encoded in protein sequences, including residue conservation, residue– residue covariation, and higher-order sequence dependencies, captures structural and functional constraints that have shaped proteins throughout evolution^3,4^. In computational protein structure prediction, effectively identifying and exploiting these evolutionary signals to improve prediction accuracy has emerged as a central challenge^5,6^.

Traditional structure prediction approaches that exploit evolutionary information infer coevolving residue pairs from multiple sequence alignments (MSAs) using statistical models and incorporate the inferred pairs as spatial restraints during structure modeling^5,7–9^. MSAs place a protein sequence in the context of evolutionarily related sequences, providing a representation of sequence variation from an evolutionary perspective. When an MSA contains a sufficient number of diverse homologous sequences, residue pairs exhibiting strong coevolutionary signals tend to be spatially proximal, thereby providing valuable restraints for structure prediction. Nevertheless, these approaches struggle to extract sufficiently reliable coevolutionary signals, particularly when evolutionary information is limited. Moreover, even when informative coevolutionary signals are available, generating accurate all-atom structures from these constraints remains challenging^5,9,10^.

Deep learning has transformed both the extraction of evolutionary information from MSAs and its integration into structure prediction^11^. MSA-based protein language models pretrained on large-scale alignments using self-supervised learning, such as MSA Transformer^12^, have substantially improved the identification of coevolving residue pairs from MSAs relative to traditional statistical approaches. Meanwhile, supervised deep-learning methods have enabled more accurate inference of spatial restraints from coevolutionary signals and other sequence-derived features^6,13,14^. Recent breakthroughs, exemplified by AlphaFold2^15^, have revolutionized the field by integrating deep-learning-based MSA representations with end-to-end protein structure prediction, enabling highly accurate modeling across a broad range of proteins^15,16^. Beyond prediction of a single conformational state, supplying AlphaFold2 with subsampled or selectively filtered MSAs containing distinct evolutionary information has enabled the modeling of alternative conformational states^17–21^. Despite these successes, existing methods still struggle to accurately predict structures for proteins with limited homologous-sequence or template information^22,23^. This reduced performance when only a few homologous sequences are available also constrains the effectiveness of MSA-sampling strategies for predicting alternative conformational states.

Single-sequence structure prediction methods, which integrate single-sequence protein language models (PLMs) with end-to-end structure prediction networks, provide a complementary approach to MSA-based methods^24–27^. Training on large-scale protein sequence datasets through self-supervised learning enables PLMs to learn protein sequence representations that implicitly encode evolutionary constraints without requiring an explicit MSA. Although single-sequence structure prediction methods generally underperform MSA-based methods such as AlphaFold2 when homologous sequences are available^23^, they can outperform MSA-based approaches for orphan proteins lacking detectable homologs^28^. Given that a single sequence can be viewed as an MSA containing only the query sequence, we therefore hypothesize that differences in generalization between single-sequence and MSA-based structure prediction models may partly arise from differences in the distribution of MSA depths (the number of homologous sequences in an MSA) during training and, consequently, in the distribution of evolutionary information available to the models. Specifically, models trained predominantly on single-sequence or shallow-MSA inputs may generalize better when homologous sequence information is limited, whereas models trained on deeper MSAs may perform better on proteins with abundant homologous sequence information. If validated, this hypothesis would imply that model generalization can be modulated by controlling the MSA-depth distributions during training, and that an appropriately designed distribution could yield models with robust performance across proteins with widely varying amounts of available evolutionary information. Previous studies^13,22,29,30^ have explored MSA sampling during training; however, they reported only modest performance gains and did not systematically investigate how the distribution of MSA depths during training influences model generalization.

A deep learning framework for all-atom protein structure prediction from MSAs was developed to investigate this hypothesis, particularly the relationship between the MSA-depth distributions encountered during training and model generalization. As illustrated in Fig. 1a (see also Methods and Supplementary Information), the framework comprises three main components: an MSA encoder, a structure module, and a confidence module. The MSA encoder extracts MSA representations and inter-residue pair representations solely from the input MSA. These representations are subsequently processed by the structure module, which predicts and iteratively refines the all-atom protein structure. The confidence module integrates structural information with MSA-derived representations to estimate per-residue LDDT-Cα scores for the predicted structure. To compel the network to rely solely on the supplied MSA for structure prediction, full-network recycling was disabled. Recycling feeds predicted structures and intermediate representations from one iteration back into the network for further refinement and is used by AlphaFold2 and other state-of-the-art protein structure prediction methods to improve prediction accuracy^31–33^. This non-recycling strategy, together with a lightweight MSA encoder, an efficient structure module and an optimized implementation, substantially reduces inference cost. On a single NVIDIA H100 PCIe GPU, the framework predicted the structure of a 1,214-residue protein from an MSA with a depth of 128 in 11.37 s, corresponding to approximately a 100-fold speedup over AlphaFold2 (version 2.3.1; weights: params_model_4) and a 10-fold speedup over AlphaFold3^31^ (version 3.0.0) (Fig. 1b).

**Figure 1:**
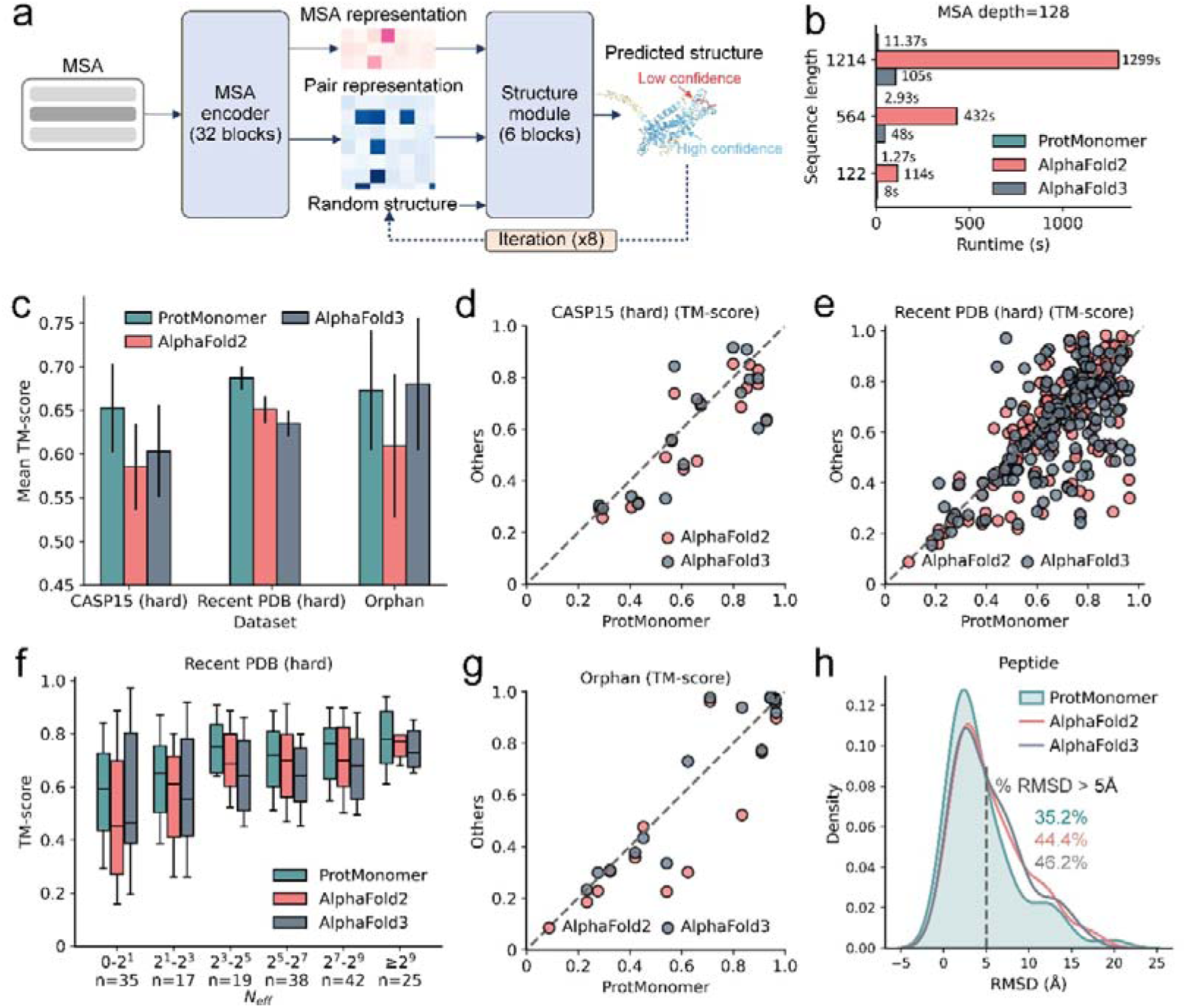
Overall comparison of ProtMonomer, AlphaFold2 and AlphaFold3 for single-conformation prediction. **a**, ProtMonomer model architecture. MSA, multiple sequence alignment. **b,** Runtime comparison of ProtMonomer and reference methods. All benchmarks were conducted on an H100-PCIe GPU. AlphaFold2 was run using a single set of model weights, and AlphaFold3 was run using a single random seed. Structural relaxation was disabled, and structural templates were not used for either AlphaFold2 or AlphaFold3. **c,** Mean TM-scores achieved by ProtMonomer, AlphaFold2 and AlphaFold3 on the CASP15 (hard), Recent PDB (hard) and Orphan datasets. The CASP15 (hard) dataset comprises 17 challenging targets, the Recent PDB (hard) dataset comprises 176 challenging proteins and the Orphan dataset comprises 15 targets. Black vertical lines indicate the standard deviation across protein targets. **d-e,** Pairwise comparisons of structures predicted by ProtMonomer (x-axis) with AlphaFold2 (light coral) and AlphaFold3 (slate grey) on CASP15 (hard) (**d**) and Recent PDB (hard) (**e**) proteins, evaluated using TM-score. **f,** Performance of ProtMonomer, AlphaFold2 and AlphaFold3 as a function of *N_eff_* on the Recent PDB (hard) dataset. Boxes indicate the interquartile range (25th–75th percentiles), center lines indicate the median and *n* denotes the number of proteins in each interval. **g,** Pairwise comparisons of structures predicted by ProtMonomer (x-axis) with AlphaFold2 (light coral) and AlphaFold3 (slate grey) on orphan proteins, evaluated using TM-score. **h,** Distribution of RMSD values for structures predicted by ProtMonomer, AlphaFold2 and AlphaFold3 across 54 peptide targets in the Peptide dataset.

Using this framework, a series of structure prediction models were trained on different MSA-depth distributions and evaluated for their ability to generalize across proteins with widely varying amounts of homologous sequence information. Comparative analysis revealed that model generalization depended strongly on the MSA-depth distribution encountered during training. Models trained on shallower MSAs performed better for proteins with limited homologous sequence information, whereas models trained on deeper MSAs performed better when richer homologous sequence information was available. Moreover, combining MSA-depth distributions integrated these complementary generalization strengths: a model trained on an equally weighted mixture of two distributions matched or exceeded the performance of either single-distribution model across proteins with a broad range of homologous sequence availability. These results suggest that exposure to a broad distribution of MSA depths during training may broaden model generalization across heterogeneous evolutionary regimes.

Based on these findings, we developed ProtMonomer, a deep learning-based method trained across a broad distribution of MSA depths to improve protein structure prediction. Benchmarking on a set of experimental structures, each with more than 5,000 homologs, showed that ProtMonomer consistently outperformed AlphaFold2 when the corresponding MSAs were downsampled across a broad range of depths. This improvement was particularly pronounced when homologous sequence information was limited or unavailable. We further evaluated ProtMonomer on single-conformation prediction across diverse benchmark settings, including CASP15 targets, recently released experimental structures, orphan proteins, and short peptides from CAMEO. Across these benchmarks, ProtMonomer performed comparably to or better than leading methods, including AlphaFold2 and AlphaFold3^31^. Beyond single-conformation prediction, we also evaluated whether these generalization improvements extend to the prediction of alternative conformational states. Approaches that combine MSA sampling with structure prediction to generate alternative conformations require models to generalize effectively from limited evolutionary information. Fold-switching proteins, which are particularly challenging for existing structure prediction methods^21,34^, provided a stringent test of this capability. Across diverse homologous sequence sampling strategies^19,35,36^, including AF-Cluster^19^, SPEACH_AF^36^, and CF-Random^35^, ProtMonomer achieved several-fold higher success rates in recovering alternative conformational states than AlphaFold2 and AlphaFold3. Moreover, confidence scores predicted by ProtMonomer provided useful guidance for identifying native-like conformations among the resulting predictions. Together, these results demonstrate that controlling the distribution of MSA depths presented during training can improve the generalization of protein structure prediction models, suggesting a general strategy for improving both single– and multiple-conformation protein structure prediction.

## Results

### Performance of ProtMonomer in single-conformation prediction

We comprehensively evaluated ProtMonomer for single-conformation prediction on four datasets spanning diverse prediction regimes: 45 publicly available CASP15 targets, 866 recently released experimentally determined structures, 15 orphan proteins with no detectable homologs in UniRef90^37^, BFD^15^, MGnify^38^ and ColabFoldDB^39^, and 54 short peptides of up to 50 residues from the CAMEO benchmark (collected between 9 April 2022 and 6 April 2024). These datasets are hereafter referred to as the CASP15, Recent PDB, Orphan, and Peptide datasets, respectively. Structures in the Recent PDB and Orphan datasets were deposited in the Protein Data Bank^40^ (PDB) between 30 March 2022 and 30 June 2024 and were filtered to exclude proteins sharing ≥40% sequence identity with any protein in the ProtMonomer training set (see Methods). For each dataset, AlphaFold2 and AlphaFold3^31^ were used as state-of-the-art comparators. ProtMonomer, AlphaFold2, and AlphaFold3 were provided with the same input MSA (see Methods for MSA generation). Although the use of structural templates has been reported to have limited effects on the performance of AlphaFold2 and AlphaFold3, templates were disabled for these methods to ensure a controlled comparison^39,41^. Prediction accuracy on the CASP15, Recent PDB, and Orphan datasets was assessed using TM-score^42^, which ranges from 0 to 1, with values ≥0.5 generally indicating a correct overall fold^43^. For the Peptide dataset, Cα root-mean-square deviation (RMSD) was used to assess structural accuracy given the short lengths of the peptides, with lower RMSD values indicating greater structural similarity to the reference structures.

ProtMonomer showed higher prediction accuracy on the CASP15 dataset, particularly for challenging targets. The Critical Assessment of protein Structure Prediction (CASP) is a community-wide blind benchmark that includes targets with limited similarity to previously characterized protein structures. Across the 45 CASP15 targets, all three methods generated high-quality predictions (TM-score ≥0.8) for a substantial proportion of targets: 68.8% for AlphaFold2, 68.8% for AlphaFold3 and 75.6% for ProtMonomer (Table S1). This high overall accuracy reduces the discriminatory power of the full benchmark. We therefore excluded proteins for which all three methods achieved a TM-score ≥0.8, leaving a challenging subset of 17 targets for further evaluation, hereafter referred to as the CASP15 (hard) dataset. On this subset, ProtMonomer achieved a mean TM-score of 0.652, compared with 0.585 for AlphaFold2 and 0.603 for AlphaFold3 (Fig. 1c). ProtMonomer also generated high-quality predictions for 35.3% of these targets, compared with 17.6% for AlphaFold2 and 17.6% for AlphaFold3. Pairwise comparisons further demonstrated the performance advantage of ProtMonomer on these challenging targets (Fig. 1d). Using a TM-score difference of 0.1 as the threshold for a substantial difference in prediction accuracy, ProtMonomer outperformed AlphaFold2 and AlphaFold3 on 41.1% and 29.4% of targets, respectively, whereas it underperformed them on 5.88% and 11.8%, respectively. Because all three methods can generate alternative predictions through MSA sampling, and AlphaFold2 can additionally produce predictions using different model weights, we performed five independent prediction runs per method and target to determine whether repeated prediction attempts improved accuracy. However, repeated prediction provided negligible improvement for all three methods (Fig. S1 and Table S2).

ProtMonomer outperformed AlphaFold2 and AlphaFold3 on challenging proteins from the Recent PDB dataset across a broad range of MSA depths. The larger Recent PDB dataset complements the relatively small CASP15 benchmark by providing broader coverage of protein structures and evolutionary contexts. As in the CASP15 analysis, all three methods generated high-quality predictions (TM-score ≥ 0.8) for a large proportion of the 866 recently released structures: 84.2% for AlphaFold2, 83.1% for AlphaFold3, and 85.2% for ProtMonomer (Table S3). After excluding proteins for which all three methods achieved a TM-score ≥0.8, 176 proteins remained as a challenging subset for further evaluation, hereafter referred to as the Recent PDB (hard) dataset. On this subset, ProtMonomer achieved a mean TM-score of 0.687, compared with 0.651 for AlphaFold2 and 0.635 for AlphaFold3 (Fig. 1c). Using TM-score ≥0.5 as the criterion for a successful prediction, ProtMonomer achieved a success rate of 86.4%, compared with 77.2% for AlphaFold2 and 74.4% for AlphaFold3 (Fig. S2 and Table S4). Pairwise comparisons showed a pattern similar to that observed for the challenging CASP15 targets (Fig. 1e). Using a TM-score difference of at least 0.1 as the threshold for a substantial performance difference, ProtMonomer outperformed AlphaFold2 on 27.0% of proteins and AlphaFold3 on 27.5%, whereas it underperformed AlphaFold2 on 10.0% and AlphaFold3 on 11.5%. Given that prediction accuracy can depend strongly on the amount of evolutionary information available, we next examined performance as a function of *N_eff_*. Relative to MSA depth, the normalized number of effective sequences (*N_eff_*) provides a more accurate measure of the information content in an MSA. The performance of the three methods was evaluated across *N_eff_* intervals, where each interval groups proteins in the Recent PDB dataset according to their *N_eff_* (Fig. 1f). ProtMonomer consistently outperformed AlphaFold2 and AlphaFold3 across these intervals (Table S5). In the lowest *N_eff_* interval (0-2), ProtMonomer achieved a median TM-score of 0.594, compared with 0.451 for AlphaFold2 and 0.465 for AlphaFold3. These results indicate that the performance advantage of ProtMonomer is maintained across varying levels of evolutionary information and is particularly pronounced when evolutionary information is highly limited.

We further evaluated the ability of ProtMonomer to predict structures for orphan proteins lacking detectable evolutionary information. Because only five orphan proteins were identified among the 866 recently released experimental structures under the original filtering criteria, we relaxed the resolution cutoff to increase the number of eligible targets (see Methods), yielding a total of 15 proteins for evaluation, hereafter referred to as the Orphan dataset. As shown in Fig. 1c, ProtMonomer achieved a mean TM-score of 0.674, outperforming AlphaFold2 (0.609) and performing comparably to AlphaFold3 (0.680). Using TM-score ≥0.5 as the criterion for a successful prediction, ProtMonomer successfully predicted 9 of 15 targets (60.0%), compared with 8 of 15 (53.3%) for AlphaFold2 and 9 of 15 (60.0%) for AlphaFold3 (Fig. 1g). Given that AlphaFold3 was trained using a substantially larger distillation dataset than ProtMonomer, comprising more than 40 million proteins, we performed an additional comparison with Boltz-1, an AlphaFold3-like reference model. Boltz-1^44^ employs an architecture and training strategy similar to those of AlphaFold3 but was trained using a distillation dataset approximately one percent the size of that used for AlphaFold3. Benchmarking showed that Boltz-1 achieved a mean TM-score of 0.641 and yielded lower TM-scores than ProtMonomer and AlphaFold3 for a substantial proportion of targets (Fig. S3). These results suggest that increasing the scale of distillation data may further improve the performance of ProtMonomer on orphan proteins.

ProtMonomer outperformed AlphaFold2 and AlphaFold3 in peptide structure prediction. Compared with larger proteins, short peptides can exhibit substantial conformational flexibility, making accurate structure prediction more challenging^45^. A total of 54 peptides collected from the Continuous Automated Model EvaluatiOn (CAMEO) benchmark^46^, including 24 structures determined by nuclear magnetic resonance (NMR) spectroscopy and 30 by X-ray crystallography, were designated as the Peptide dataset for model evaluation. ProtMonomer achieved a mean RMSD of 4.63 Å on the Peptide dataset, compared with 5.55 Å for AlphaFold2 and 5.44 Å for AlphaFold3. Consistently, the RMSD distributions showed that structures predicted by ProtMonomer had overall lower RMSD values than those predicted by AlphaFold2 and AlphaFold3 (Fig. 1h; pairwise comparisons are shown in Fig. S4). Using an RMSD threshold of less than 5 Å as the criterion for a successful prediction, ProtMonomer achieved a success rate of 64.8%, compared with 55.5% for AlphaFold2 and 53.7% for AlphaFold3. We further grouped the peptides according to their *N_eff_* values and evaluated the three methods across different *N_eff_* intervals. Although the three methods showed comparable performance in the lowest *N_eff_* interval, ProtMonomer benefited more substantially from increasing MSA depth. In contrast, AlphaFold2 and AlphaFold3 showed more limited improvements with additional evolutionary information (Fig. S5 and Table S6).

### Accurate prediction of fold-switching protein conformations

The fold-switching protein dataset assembled by Chakravarty et al.^34^ was used to assess the ability of ProtMonomer to predict alternative conformational states. To our knowledge, this dataset represents the most comprehensive collection of experimentally characterized fold-switching proteins currently available. The original dataset contained 92 pairs of experimentally determined fold-switching proteins collected from the PDB, with each pair comprising identical or nearly identical sequences that adopt distinct secondary and tertiary structural arrangements. For our evaluation, 68 pairs with identical sequences were retained.

Motivated by the hypothesis that MSAs encode information about protein structural heterogeneity, a general approach for predicting alternative protein conformations is to partition a full MSA into multiple sub-MSAs containing distinct evolutionary information and use these sub-MSAs as inputs to structure prediction models^47^. Several MSA sampling strategies based on different principles have been proposed, including clustering sequences according to sequence similarity, *in silico* mutagenesis, and random subsampling. However, these sampling methods often generate relatively shallow sub-MSAs, and the reduced prediction accuracy of structure prediction models under limited evolutionary information may constrain their ability to recover alternative conformational states. In this study, we combined three MSA sampling methods, AF-Cluster^19^ (based on sequence clustering), SPEACH_AF^36^ (based on *in silico* mutagenesis) and CF-Random^35^ (based on random subsampling), with ProtMonomer, AlphaFold2 and AlphaFold3 to predict alternative conformations for each fold-switching protein. Structural templates were disabled for AlphaFold2 and AlphaFold3 to enable a controlled comparison. Given the relatively short sequence lengths of the fold-switching proteins (Fig. S6), we used RMSD to assess structural similarity between the predicted structures and the experimentally determined conformations.

ProtMonomer achieved substantially higher success rates in predicting both conformations than AlphaFold2 and AlphaFold3 across diverse MSA sampling strategies. Given the inherent difficulty of multiple-conformation prediction, we used a relatively lenient RMSD threshold of <5 Å to define structural similarity. A method was considered successful for a given protein if the resulting structural ensemble contained at least one model within 5 Å RMSD of each of the two experimentally determined conformations. As shown in Fig. 2a, under the AF-Cluster sampling strategy, ProtMonomer achieved a success rate of 54.4%, compared with 14.7% for AlphaFold2 and 13.2% for AlphaFold3. Notably, the limited performance of AlphaFold2 and AlphaFold3 was consistent with the findings of Chakravarty et al., who previously reported challenges in recovering alternative conformations for both methods on this dataset. Although success rates were lower under the CF-Random and SPEACH_AF sampling strategies, the overall advantage of ProtMonomer was maintained. Under CF-Random, ProtMonomer achieved a success rate of 41.1%, compared with 10.3% for AlphaFold2 and 11.7% for AlphaFold3. Under SPEACH_AF, the corresponding success rates were 14.7%, 5.8%, and 0%, respectively. These results demonstrate that ProtMonomer substantially improves the prediction of alternative conformations and consistently enhances the effectiveness of different MSA sampling strategies.

**Figure 2:**
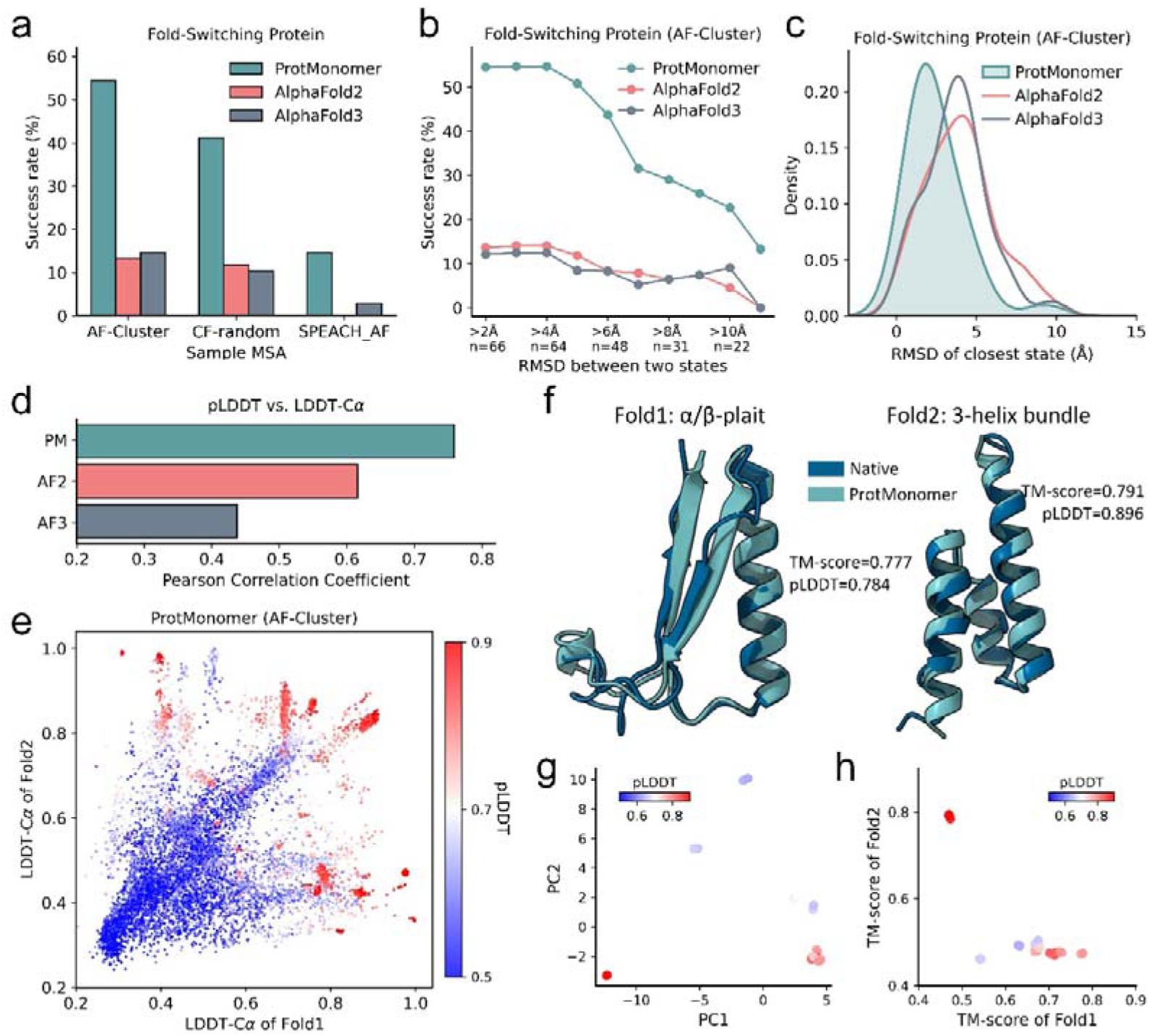
Performance of ProtMonomer, AlphaFold2, and AlphaFold3 on fold-switching proteins. **a**, Success rates of ProtMonomer, AlphaFold2, and AlphaFold3 for recovering both experimentally determined conformations across diverse MSA sampling strategies. **b,** Success rates of ProtMonomer, AlphaFold2, and AlphaFold3 for recovering both experimentally determined conformations for fold-switching proteins stratified by the magnitude of the structural difference between their two experimental conformations using the AF-Cluster sampling strategy. *n* denotes the number of proteins in each interval. **c,** Distributions of the minimum RMSDs of structures predicted by ProtMonomer, AlphaFold2, and AlphaFold3 relative to the closest experimentally determined conformation. **d,** Pearson correlation coefficients (PCCs) between model-predicted LDDT-Cα (pLDDT) and LDDT-Cα for structures predicted for the fold-switching dataset. For each prediction, LDDT-Cα was calculated against both experimentally determined conformations, and the higher of the two values was used as the measure of structural accuracy. AF2, AlphaFold2; AF3, AlphaFold3. **e,** Comparison of LDDT-Cα values of structures predicted by ProtMonomer using AF-Cluster relative to the Fold1 (x-axis) and Fold2 (y-axis) experimental conformations for the fold-switching dataset, colored by pLDDT. **f,** The Sa1 protein switches reversibly between α/β-plait (PDB ID: 8e6y, Fold1) and 3α-helix (PDB ID: 2fs1, Fold2) folds in response to temperature changes. ProtMonomer accurately recovered both experimental conformations. **g,** Principal component analysis of Cα-Cα distance maps derived from structures predicted by ProtMonomer, colored by pLDDT. **h,** TM-scores of 24 ProtMonomer predictions relative to the Fold1 and Fold2 conformations of the Sa1 protein, showing that the predicted structural ensemble captures both the α/β-plait and 3α-helix conformations.

We further investigated the relationship between the ability of the three methods to recover both conformations and the structural difference between the two experimentally determined conformations of fold-switching proteins. The AF-Cluster strategy was adopted for MSA sampling because it consistently outperformed CF-Random and SPEACH_AF across all three structure prediction methods. Proteins were grouped according to the magnitude of the structural difference between their two experimental conformations, and the success rate for recovering both conformations was evaluated within each group for each method. As shown in Fig. 2b, ProtMonomer consistently outperformed AlphaFold2 and AlphaFold3 across all groups (Table S7), indicating that its performance advantage was maintained across a broad range of conformational changes. Notably, the success rates of all three methods decreased markedly as the structural difference increased, suggesting that recovering both conformations becomes increasingly challenging for fold-switching proteins undergoing larger conformational changes. However, even for proteins with structural differences exceeding 11 Å, the success rates of both AlphaFold2 and AlphaFold3 dropped to 0%, whereas ProtMonomer retained a success rate of 13.3%, highlighting the greater robustness of ProtMonomer.

Given the challenges that conformational flexibility poses for structure prediction, we also investigated the ability of ProtMonomer to predict at least one experimentally observed conformation for each fold-switching protein. For each protein and prediction method, the minimum RMSD across all predicted structures relative to either of the two experimentally determined conformations was used to assess prediction accuracy. As shown by the distribution of minimum RMSD values across the dataset (Fig. 2c), ProtMonomer outperformed AlphaFold2 and AlphaFold3 in recovering at least one experimentally observed conformation. Specifically, ProtMonomer achieved a minimum RMSD <5 Å for 92.6% of proteins, compared with 72.1% for AlphaFold2 and 82.4% for AlphaFold3. Using a stricter RMSD threshold of <2 Å to define high-quality predictions, ProtMonomer achieved this criterion for 45.5% of proteins, compared with 19.1% for AlphaFold2 and 20.6% for AlphaFold3.

After establishing the ability of ProtMonomer to predict multiple conformations, a key question was whether native-like conformations could be reliably identified within the predicted structural ensemble. Given that all three structure prediction methods provide predicted LDDT-Cα (pLDDT) scores as estimates of prediction confidence, we evaluated how well these scores distinguish native-like conformations among the generated structures. The Local Distance Difference Test (LDDT-Cα)^48^ quantifies structural similarity between two structures, with higher values indicating greater structural similarity. Because each fold-switching protein has two experimentally determined conformations and a predicted structure may resemble either conformation, we calculated the LDDT-Cα of each prediction against both experimental structures and used the higher value as the measure of structural accuracy. Fig. 2d shows the Pearson correlation coefficients (PCCs) between pLDDT and LDDT-Cα for structures predicted by the three methods across the fold-switching dataset. ProtMonomer exhibited a PCC of 0.759, substantially higher than those of AlphaFold2 (0.616) and AlphaFold3 (0.437), indicating that its pLDDT scores provide more reliable estimates of structural accuracy. Head-to-head comparisons further showed that AlphaFold3 tended to overestimate the quality of its predicted structures (Fig. S7). Fig. 2e shows the LDDT-Cα values of structures predicted by ProtMonomer relative to the two experimentally determined conformations, with corresponding results for AlphaFold2 and AlphaFold3 shown in Fig. S8. Predicted structures with pLDDT >0.8 generally exhibited high structural similarity (LDDT-Cα >0.8) to at least one of the experimentally determined conformations, suggesting that pLDDT can be used to identify native-like conformations within structural ensembles generated by ProtMonomer.

The ability of ProtMonomer to predict multiple conformations was further examined using the Sa1 protein^49^, which exhibits temperature-dependent reversible switching between a 3α-helix bundle and an α/β-plait fold. Chakravarty et al.^34^ showed that, despite generating thousands of structures with AlphaFold2 and AlphaFold3 using various subsampled MSAs, both methods reproduced only the α/β-plait fold. As an initial test, we sampled four sub-MSAs at each of six MSA depths (2, 4, 8, 16, 32 and 64) and generated one structure from each sampled MSA using ProtMonomer. Principal component analysis^50^ of the 24 predicted structures revealed that high-confidence predictions (pLDDT >0.8) clustered into two groups (Fig. 2g), corresponding to two distinct structural conformations. Structural similarity analysis against the two experimentally determined conformations (Fig. 2h) further confirmed that the two clusters corresponded to the two experimental states. Overall, 4 of the 24 predicted structures matched the 3α-helix conformation (TM-score >0.5), whereas the remaining 20 matched the α/β-plait fold.

### Investigating the generalization of protein structur**e** prediction models

To investigate how the generalization capability of protein structure prediction models depends on the MSA-depth distribution used during training, a comprehensive collection of 16,225 high-resolution experimentally determined protein structures, each with an MSA containing more than 5,000 sequences, was assembled as the training dataset (see Methods). For a given MSA-depth distribution, the MSA of each protein was sampled according to the prescribed distribution before being used for model training. This procedure ensured that the MSA-depth distribution encountered by the models during training matched the prescribed distribution. In addition, we curated an evaluation dataset comprising 532 proteins that were non-redundant with the training dataset, each with a high-resolution experimentally determined structure and an MSA containing more than 5,000 sequences (see Methods). The MSAs for these proteins were downsampled to depths ranging from 1 to 8,192 to comprehensively evaluate model performance across MSA depths^22^.

Because any MSA-depth distribution can be represented as a weighted combination of delta distributions centered at discrete MSA depths, we initially investigated the generalization ability of protein structure prediction models trained on MSAs of fixed depths. Specifically, we trained three models using MSAs downsampled to depths of 32, 128 and 512 (see Methods for model training) and evaluated each model on the evaluation dataset using MSAs downsampled to a range of depths. As shown in Fig. 3a-b, the models exhibited complementary generalization capabilities: models trained with shallower MSAs achieved higher prediction accuracy than models trained with deeper MSAs when evaluated with shallow MSAs but showed lower accuracy when evaluated with deeper MSAs. These results demonstrate that the generalization capabilities of protein structure prediction models can be modulated by adjusting the MSA-depth distribution encountered during training. Notably, for a given model, once the MSA-depth during inference exceeded its training depth, further increases in MSA depth provided little additional improvement (Fig. 3a). This saturation suggests that the ability of a model to exploit additional evolutionary information is constrained by the range of MSA depths encountered during training.

**Figure 3:**
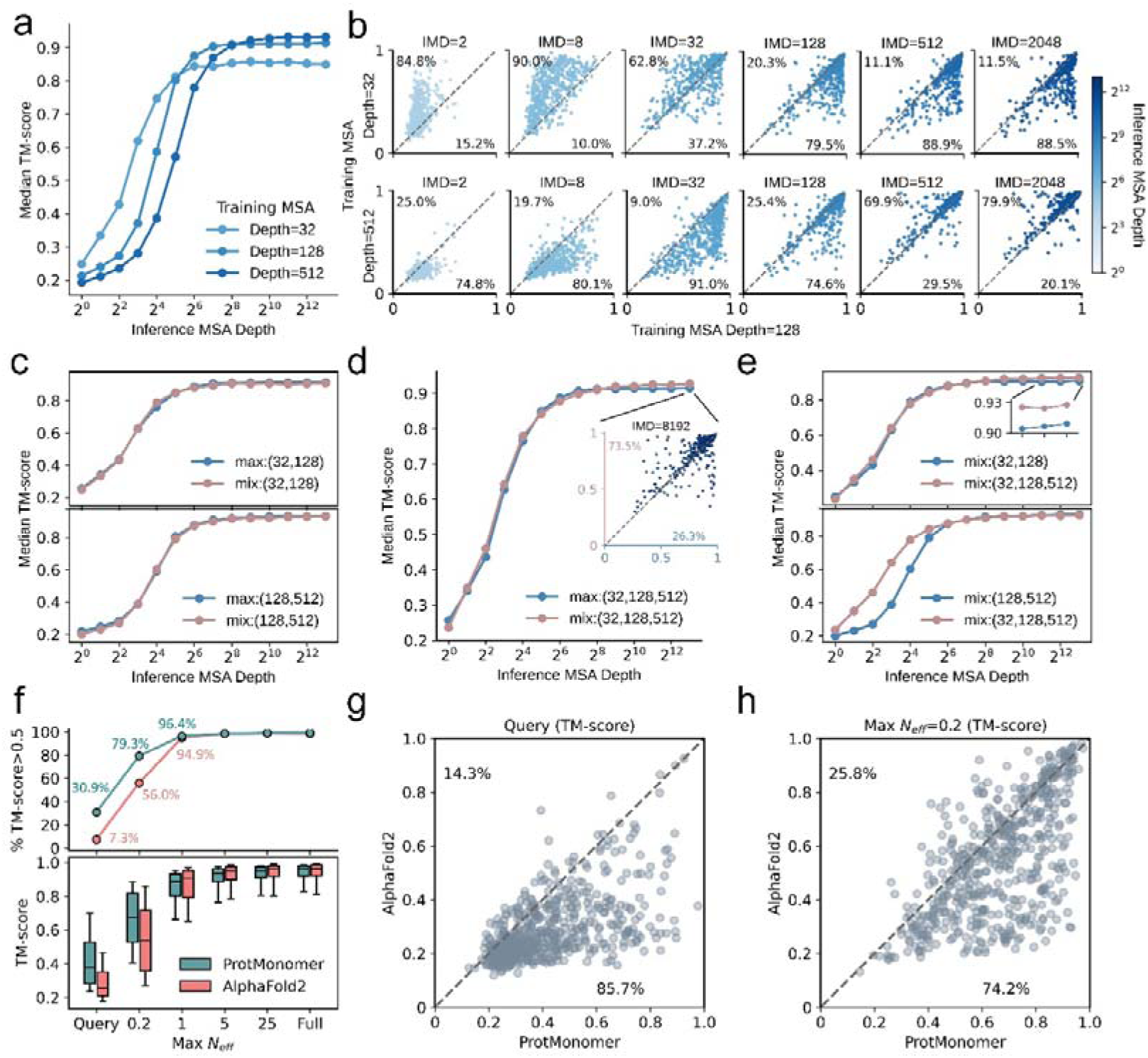
Generalization of protein structure prediction models trained on different MSA-depth distributions. **a**, Median TM-scores of predictions from models trained at MSA depths 32, 128, and 512 on the evaluation dataset, showing distinct generalization performance across inference MSA depths. **b,** Pairwise comparisons of models trained at MSA depths of 32, 128 and 512, evaluated across varying inference MSA depths (IMD) on the evaluation dataset. Percentages indicate the fractions of proteins falling within the upper or lower triangular regions of the plot. **c,** The upper panel shows that the model trained on a mixture of depths 32 and 128 (mix:(32,128)) achieves performance comparable to the best of the models trained separately at depths 32 or 128 (max:(32,128)) on the evaluation dataset. The lower panel shows a similar trend: the performance of the model trained on a mixture of depths 128 and 512 (mix:(128,512)) matches the best performance of models trained separately at depths 128 or 512 (max:(128,512)). **d,** Performance of the model trained on the mixture of MSA depths of 32, 128 and 512 (denoted as mix:(32,128,512)) matches or exceeds the best performance of independently trained models at MSA depths 32, 128 and 512 (denoted as max:(32,128,512)). **e,** Performance comparison between the model trained on a mixture of depths 32, 128 and the model trained on a mixture of depths 32, 128, and 512 (upper panel). The lower panel shows the performance comparison between model trained on a mixture of depths 128, 512 and model trained on a mixture of depths 32, 128, 512. **f,** TM-score distributions (lower panel) and success rates (upper panel) of ProtMonomer and AlphaFold2 on the evaluation dataset across different inference *N_eff_*. Boxes indicate the interquartile range (25th–75th percentiles) and center lines indicate the median. **g-h,** Pairwise comparison of ProtMonomer (x-axis) and AlphaFold2 (y-axis) predictions on the evaluation dataset with different MSA depth constraints (**g:** MSA depth equal to 1; **h:** *N_eff_* of each sampled MSA equal to 0.2).

Given that delta distributions centered at different MSA depths can serve as fundamental components for developing protein structure prediction models, exploring the impact of combining these components in different ways on the generalization capabilities of the resulting models is of particular importance. The simplest case involves combining these components pairwise with equal weights. We developed two types of models to investigate this. The first was trained on an equal mixture of delta distributions centered at MSA depths a and b, denoted as mix:(a,b), where a and b represent the centers of the delta distributions. The second was a joint model comprising two models trained separately at depths a and b, for which the prediction with the higher TM-score to the experimental structure was selected retrospectively, denoted as max:(a,b). Specifically, we developed four models: mix:(32,128), mix:(128,512) and their corresponding joint models, max:(32,128) and max:(128,512). Fig. 3c illustrates the performance of these models on the evaluation dataset where the MSAs were downsampled at various depths. Across evaluation MSA depths, the mixed-distribution models achieved performance comparable to or better than that of the corresponding joint models. These results indicate that training on an equal mixture of MSA-depth distributions can integrate the generalization capabilities associated with individual training depths within a single model. We further evaluated whether this behavior extended to a mixture of three MSA depths by comparing mix:(32,128,512) with its corresponding joint model, max:(32,128,512) (Fig. 3d). The mix:(32,128,512) model achieved performance comparable to or better than that of max:(32,128,512) across evaluation MSA depths, further supporting the conclusion that a single model trained on a mixture of MSA depths can integrate the generalization capabilities of models trained separately at individual depths.

We subsequently explored which MSA-depth distribution may maximize the generalization capacity of protein structure prediction models. Considering that a complex MSA-depth distribution can be extended by introducing an additional delta component centered at an MSA depth outside its current support, we asked whether placing the delta component below the lower bound or above the upper bound of the complex distribution results in greater improvements in model performance. We compared the performance of the model mix:(32,128,512) with those of mix:(32,128) and mix:(128,512) to investigate this (Fig. 3e). Comparing mix:(32,128) and mix:(32,128,512) reveals that, based on the distribution equally combining delta distributions centered at MSA depths of 32 and 128, increasing the sampling probability in the deeper region leads to only modest performance improvement. In contrast, comparing mix:(128,512) and mix:(32,128,512) showed that increasing the sampling probability in the shallow-MSA regime based on the distribution equally combining delta distributions centered at MSA depths of 128 and 512, produced substantial performance gains in this regime. These results suggest that increasing the representation of shallow MSAs during training is more beneficial for improving model generalization. Notably, predictions for proteins evaluated with deeper MSAs (depth ≥256) were already highly accurate, with median TM-scores ≥0.9, whereas predictions for proteins evaluated with shallower MSAs remained substantially less accurate and therefore offered greater scope for improvement.

Drawing on these insights, we designed a multi-scale MSA-depth sampling distribution for ProtMonomer training, spanning 1–2,048 homologous sequences. The distribution consisted of four uniform sampling intervals, 1–32, 32–128, 128–512, and 512–2,048 homologous sequences, with equal sampling weights. In parallel, additional experimentally determined and distilled structures were incorporated into the training dataset (see Methods). After the initial training stage had converged, two additional training stages, which used modified MSA-depth distributions with lower sampling probabilities for deep MSAs and higher probabilities for shallow MSAs, further improved performance in the low-MSA-depth regime (Table S8 and Fig. S9). We further benchmarked the resulting ProtMonomer against AlphaFold2 on the evaluation dataset using MSAs downsampled across a range of *N_eff_* values. As shown in Fig. 3f–h, ProtMonomer outperformed AlphaFold2 across varying *N_eff_* values. The improvement was particularly pronounced when only the single query sequence (Query) (Fig. 3g) or a small number of homologous sequences (*N_eff_* = 0.2) (Fig. 3h) were available for prediction. Under these conditions, ProtMonomer achieved success rates (TM-score ≥0.5) of 30.9% and 79.3%, respectively, corresponding to improvements of 23.6 and 23.3 percentage points over AlphaFold2.

## Discussion

Although deep learning-based protein structure prediction methods have achieved remarkable success in single-conformation prediction, substantial challenges remain for short peptides, proteins with limited evolutionary or template information, and proteins that adopt multiple conformational states. Motivated by the different generalization behaviors of MSA-based and PLM-based structure prediction methods, we investigated how the MSA-depth distribution encountered during training influences model generalization. Our analyses show that models trained on MSA-depth distributions representing different amounts of available evolutionary information exhibit complementary generalization capabilities, and that training on mixtures of these distributions can combine their complementary strengths within a single model. Building on these findings, we developed ProtMonomer, a deep learning-based structure prediction method trained across a broad distribution of MSA depths to improve generalization across different levels of evolutionary information.

ProtMonomer improved both single-conformation prediction and the recovery of alternative protein conformations. Comprehensive benchmarking of single-conformation prediction across CASP15 targets, recently released experimental structures, orphan proteins, and short peptides showed that ProtMonomer performed comparably to or better than AlphaFold2 and AlphaFold3, with particularly strong performance on challenging targets. Notably, despite these improvements, prediction accuracy for short peptides and orphan proteins remains limited. Integrating ProtMonomer into established workflows such as MassiveFold^51^ and DeepMSA2^52^ may further improve performance on these challenging targets. For fold-switching proteins, ProtMonomer substantially improved the recovery of alternative experimentally observed conformations across diverse MSA sampling strategies compared with AlphaFold2 and AlphaFold3, while its pLDDT scores provided more reliable estimates for identifying native-like structures within conformational ensembles. In addition, ProtMonomer achieved substantially faster inference than AlphaFold2 and AlphaFold3, facilitating large-scale protein structure prediction.

ProtMonomer was developed for monomeric protein structure prediction, but the relationship between training-time MSA-depth distributions and model generalization may extend to other MSA-based biomolecular structure prediction tasks. Future work could investigate whether analogous strategies for controlling training-time MSA-depth distributions improve generalization in other prediction tasks, including protein complex and RNA structure prediction.

## Methods

### Model architecture

#### Main framework

The MSA encoder consists of 32 blocks based on a slightly modified Evoformer^15^ architecture and is designed to extract informative MSA and inter-residue pair representations (see Supplementary Information for details). The structure module comprises six equivariant structure blocks that iteratively predict and refine the all-atom protein structure based on these representations. The confidence module comprises three blocks with an architecture similar to that of the MSA encoder and predicts the LDDT-Cα scores of the predicted structures by integrating structural information with MSA-derived representations.

#### Structure block

Each structure block consists of four modules: MSA2Atom, InterResidueAttn, IntraResidueAttn, and a feed-forward network (FFN) module (see Supplementary Information for details). The block takes as input the MSA and pair representations generated by the MSA encoder, the predicted structure from the previous iteration, and the current structure representation. The structure is randomly initialized^53^, whereas the initial structure representation is derived solely from the query sequence. The MSA representations are randomly subsampled and processed by the MSA2Atom module. The resulting MSA representations, together with the pair representations, predicted structure, and current structure representation, are then integrated by the InterResidueAttn module to update the structure representation based on inter-residue interactions. The IntraResidueAttn module subsequently refines the structure representation using intra-residue information, taking both the current structure representation and the predicted structure as inputs. Finally, the FFN module further transforms the structure representation in a channel-wise manner.

### Training and validation datasets

We retrieved all protein structures deposited in the Protein Data Bank (PDB) before 30 December 2021, together with AlphaFold2-predicted monomeric structures from the OpenProteinSet^54^ (approximately 270,000 predicted structures) and the PSP dataset^55^ (approximately 760,000 predicted structures), to construct the ProtMonomer training datasets. Experimental monomeric structures from the PDB were filtered according to the following criteria: structures were excluded if they (1) had a reported resolution worse than 9 Å, (2) contained non-standard residues, or (3) contained fewer than five resolved residues. After filtering, the remaining structures were clustered at 100% sequence identity, and one representative structure was randomly selected from each cluster. This procedure yielded 132,905 protein structures, collectively referred to as the experimental dataset and used as the primary training dataset for ProtMonomer. Proteins in the experimental dataset were further clustered at 40% sequence identity using CD-HIT, resulting in 32,009 clusters.

The large collection of AlphaFold2-predicted structures was used as additional training data after removing sequences redundant with any experimental structures, including those in the experimental, validation, and test datasets. Specifically, the predicted structures were clustered together with all experimental structures from these datasets using CD-HIT^56^ at 40% sequence identity with the parameters –c 0.4 –M 0 –T 0 –g 1 –n 2 –G 0 –aS 0.8. Predicted structures belonging to clusters containing any experimental structure were removed. This procedure yielded 905,643 predicted structures, collectively referred to as the distillation dataset.

Monomers deposited in the PDB between 30 December 2021 and 30 March 2022 were used to construct the validation dataset. After applying the same filtering criteria used for the experimental dataset, the remaining candidate validation proteins were clustered together with proteins from the experimental dataset using CD-HIT at 40% sequence identity to eliminate redundancy. Candidate proteins belonging to clusters containing any protein from the experimental dataset were excluded to minimize redundancy. Representative proteins were then retained from clusters containing exclusively candidate validation proteins, resulting in a validation dataset comprising 368 proteins.

In addition, we constructed a high-quality experimental dataset to investigate the generalization capabilities of protein structure prediction models. Proteins from the experimental dataset were further filtered according to the following criteria: (1) a reported resolution of no worse than 2.5 Å; and (2) more than 5,000 homologous sequences in the corresponding MSA. This filtering yielded 56,116 proteins. These proteins were subsequently clustered at 40% sequence identity using CD-HIT, and one representative protein was retained from each cluster. The resulting high-quality experimental dataset comprised 16,225 proteins.

### Test datasets

#### CASP15

All 45 publicly available proteins from https://predictioncenter.org/download_area/CASP15/targets/ were collected to constitute the CASP15 test dataset. Because the ProtMonomer experimental training set was restricted to PDB structures deposited before 30 December 2021, whereas CASP15 targets were released beginning in May 2022, the CASP15 targets postdated the structural training cutoff.

#### Recent PDB dataset and self-evaluation dataset

We collected experimental protein structures from the PDB to develop these two benchmark datasets using the following criteria: (1) deposition date between 30 March 2022 and 30 June 2024; (2) reported resolution better than 2.5 Å; (3) consisting solely of standard amino acid residues; and (4) containing at least five fully resolved residues. The remaining structures were then processed using two separate procedures to generate the Recent PDB dataset and the self-evaluation dataset.

To construct the Recent PDB dataset, we removed structures redundant with those in the experimental and validation datasets. Specifically, candidate benchmark proteins were clustered together with proteins from the experimental and validation datasets using CD-HIT at 40% sequence identity. Representative proteins were retained only from clusters containing exclusively candidate benchmark proteins. This procedure yielded a Recent PDB dataset comprising 866 protein structures.

To construct the self-evaluation dataset, we first removed proteins redundant with the experimental and validation datasets using the same procedure described above. We then further required each protein to have an MSA containing more than 5,000 homologous sequences. This procedure yielded a self-evaluation dataset comprising 532 protein structures.

#### Peptide

We collected all proteins included in the Continuous Automated Model EvaluatiOn (CAMEO) benchmark between 9 April 2022 and 6 April 2024 (https://cameo3d.org). From these targets, we selected the 54 peptides with sequence lengths of up to 50 residues to constitute the Peptide dataset.

#### Orphan protein

We collected protein structures deposited in the PDB using criteria similar to those used to construct the Recent PDB dataset, except that the resolution cutoff was relaxed to 9 Å. Among these structures, 15 proteins for which no homologous sequences could be identified using the search procedure described in the “MSA generation” section were retained to constitute the Orphan dataset.

#### Fold-switching protein

Chakravarty et al.^34^ curated a dataset comprising 92 pairs of fold-switching proteins. Each pair contains two experimentally determined structures corresponding to nearly identical protein sequences. From this dataset, we retained 68 pairs for which the two structures had identical amino-acid sequences for further analysis.

### MSA generation

For each protein with an experimentally determined structure, we generated four MSAs, removed duplicate sequences at 100% sequence identity, and used the resulting alignments as MSA inputs. Two MSAs were generated by searching the UniRef90^37^ database (downloaded on 17 July 2024) and the MGnify^38^ database (downloaded on 12 July 2024) using JackHMMER^57^ (version 3.4) with parameters consistent with AlphaFold3: –N 1 –E 0.0001 –incE 0.0001 –F1 0.0005 –F2 0.00005 –F3 0.0000005. A third MSA was generated by searching the ColabFoldDB sequence database (version 202108) using the colabfold_search tool with default parameters^39^. Finally, a fourth MSA was generated by searching the BFD^15^ database using HHblits^58^ (version 3.3) with parameters consistent with AlphaFold3: –n 3 –e 0.001 – realign_max 100000 –maxfilt 100000 –min_prefilter_hits 1000 –p 20 –Z 500.

For proteins with predicted structures, we used their corresponding MSAs provided by the OpenProteinSet or PSP dataset as complete sequence alignments.

### Training protocol

ProtMonomer was trained in three stages with progressively modified MSA-depth sampling distributions and training settings (Table S8). Proteins were sampled from the experimental and distillation datasets using stage-specific probabilities. In stage 1, proteins were sampled from the experimental and distillation datasets with probabilities of 0.25 and 0.75, respectively. The distillation probability was reduced to 0.5 in stage 2, and distilled structures were excluded in stage 3^44^. For proteins in the distillation dataset, residues with pLDDT < 0.8 were masked. Proteins in the distillation dataset were sampled with equal probability, whereas proteins in the experimental dataset were sampled with probability proportional to 1/*N_cluster_* × *min* (*max*(L, 128), 512), where *N_cluster_* is the size of the corresponding sequence cluster and L is the protein sequence length.

In stage 1, the sampling distribution comprised four intervals spanning 1–32, 32– 128, 128–512, and 512–2,048 homologous sequences. Each interval was assigned a sampling probability of 0.25, and the MSA depth was sampled uniformly within the selected interval. For each protein selected for training, an MSA depth *N_sample_* was sampled from this distribution and used as the maximum number of homologous sequences included during training. To limit GPU memory consumption and reduce training cost, the total number of residues in an MSA was constrained to at most 256×1280. Accordingly, the final MSA depth was defined as *N_final_* = *min* (*N_sample_*, *N_orig_*, [256×1280/*L*]), where *N_orig_* is the number of homologs in the original MSA and L is the sequence length. Protein sequences were cropped to a maximum length of 256 residues during this stage. For each selected training protein, 15% of residues in the sampled MSA were selected for masked-token prediction. Of these selected residues, 80% were replaced with a special mask token, 10% were replaced with a randomly selected amino acid, and 10% were left unchanged.

In stage 2, the MSA-depth distribution comprised five intervals: (1,8), (8,64), (64,512), (512,2048), and (2048,16384), each with a sampling probability of 0.2. The maximum MSA input size was increased to 340×8192, and protein sequences were cropped to a maximum length of 768 residues.

In stage 3, the MSA-depth distribution was shifted toward shallower MSAs using seven intervals: (1,4), (4,16), (16,64), (64,256), (256,1024), (1024,4096), and (4096,16384), with corresponding sampling probabilities of 0.3, 0.2, 0.1, 0.1, 0.1, 0.1, and 0.1. The maximum MSA input size and sequence crop length remained 340×8192 and 768 residues, respectively.

Similar to AlphaFold2, we defined loss functions for individual network modules. The structure module was supervised using the all-atom FAPE loss, whereas the MSA encoder was supervised using the masked amino-acid prediction loss and distogram loss. The confidence module was supervised using the pLDDT and pAE losses. The corresponding loss weights were set to 2, 1, 0.3, 0.01, and 0.01, respectively. In addition, the final loss for each training example was multiplied by the square root of the number of residues remaining after cropping. In stage 3, we replaced the all-atom FAPE loss with an all-atom and all-frame FAPE loss (Supplementary Information, Training Loss) to reduce steric clashes. We also excluded distilled structures from the training data during this stage. Whereas the all-atom FAPE loss primarily focuses on learning atom positions relative to Cα atoms, the all-atom and all-frame FAPE loss assigns equal weight to relative positions between all atom pairs. Compared with the stage 2 model, the stage 3 model reduced the average number of clashing atom pairs from approximately 5,000 to approximately 200 while maintaining nearly identical predictive performance.

ProtMonomer and all ablation models were implemented in PyTorch and trained on NVIDIA H100 and A100 GPUs. To accommodate the high memory requirements, we employed FlashAttention^59^, mixed-precision training with bfloat16, gradient checkpointing for each MSA encoder and structure-module layer, and gradient accumulation. Model parameters were optimized using AdamW^60^ with a weight decay of 0.01 and stage-specific learning rates of 0.001, 0.0005, and 0.0001 for stages 1, 2, and 3, respectively. To further improve training stability, per-sample gradients were clipped to a global norm of 0.1. ProtMonomer was trained with a batch size of 24. For models used to investigate generalization capabilities, the batch size was reduced to 4, and the models were trained using the stage 1 protocol for 25 epochs on the high-quality experimental dataset until near convergence (Fig. S10).

### Evaluation metrics

The similarity between predicted and experimental structures was assessed using the LDDT-Cα, TM-score and RMSD metrics. In addition, MSA information content was quantified using the normalized effective number of sequences, *N_eff_*.

The LDDT-Cα between predicted and experimental structures was calculated using the LDDT package (https://swissmodel.expasy.org/lddt). TM-score and RMSD were calculated using the TM-score package (https://zhanggroup.org/TM-score/). The *N_eff_*. for each MSA was calculated as

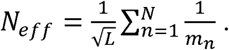

where L is the number of columns of the MSA, N is the total number of sequences in the MSA, 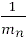 is the weight of the n-th sequence with *m_n_* being the number of sequences in the MSA that share >80% sequence identity with the n-th sequence.

### Benchmark methods

AlphaFold3 was obtained from its GitHub repository (https://github.com/google-deepmind/alphafold3). AlphaFold2 was obtained from its GitHub repository (https://github.com/google-deepmind/alphafold). Boltz-1 was obtained from its GitHub repository (https://github.com/jwohlwend/boltz). AF-Cluster was obtained from its GitHub repository (https://github.com/HWaymentSteele/AF_Cluster). CF-Random was obtained from its GitHub repository (https://github.com/ncbi/CF-random_software). SPEACH_AF was obtained from its GitHub repository (https://github.com/RSvan/SPEACH_AF).

## Acknowledgements

We thank Prof. Xin Liu at Shanghai Jiao Tong University for contributing to the conceptualization of the manuscript. We thank the developers and contributors of open-source tools and datasets including AlphaFold, OpenFold, PDB, OpenProteinSet, PSP dataset and others for greatly accelerating our research. This work was supported by National Key R&D Program of China (2022YFA1004800, 2025YFF1207900), Natural Science Foundation of China (T2350003, T2341007, 12131020, 42450084, 42450135, 12326614, 12404236 and 12426310), Zhejiang Province Vanguard Goose-Leading Initiative (2025C01114), Science and Technology Commission of Shanghai Municipality (23JS1401300), and Hangzhou Institute for advanced study of UCAS (2024HIAS-P004), and JST Moonshot R&D (JPMJMS2021).

## Author contributions

L.C. and Y.S. conceived the study. Y.S. developed ProtMonomer and performed the experiments. Y.S. analyzed the results and wrote the manuscript. L.C. and S.Z. contributed to data and method visualization. S.Z. curated the fold-switching protein dataset. L.C. supervised the research and provided funding. All authors discussed the results and commented on the manuscript.

## Data and code availability

The source code and pretrained weights for ProtMonomer are publicly available at https://github.com/yunda-si/ProtMonomer.

## Supplementary Information

### Figures

**Figure S1:**
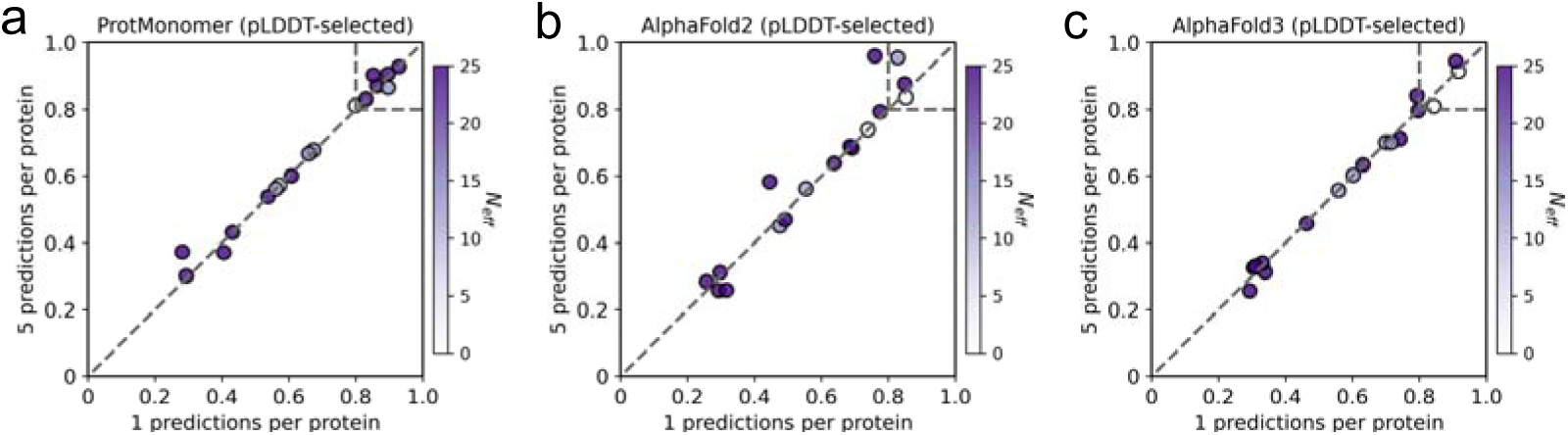
Performance of ProtMonomer. (**a**), AlphaFold2 (**b**) and AlphaFold3 (**c**) on challenging CASP15 targets with five predictions generated per protein. The prediction with the highest pLDDT score was selected as the final prediction and evaluated using TM-score.

**Figure S2:**
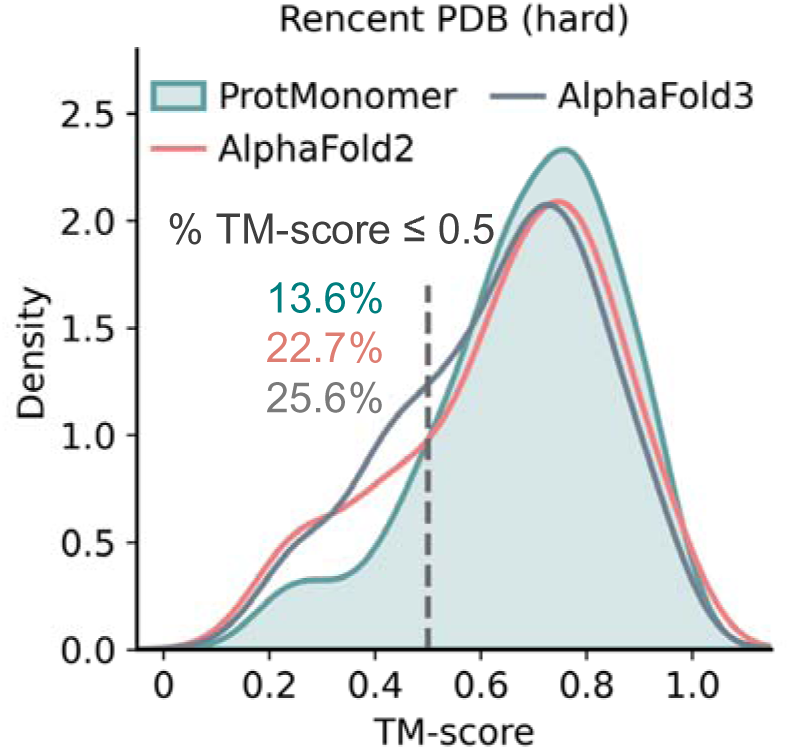
TM-score distributions of ProtMonomer, AlphaFold2 and AlphaFold3 across 176 challenging proteins from the Recent PDB dataset.

**Figure S3:**
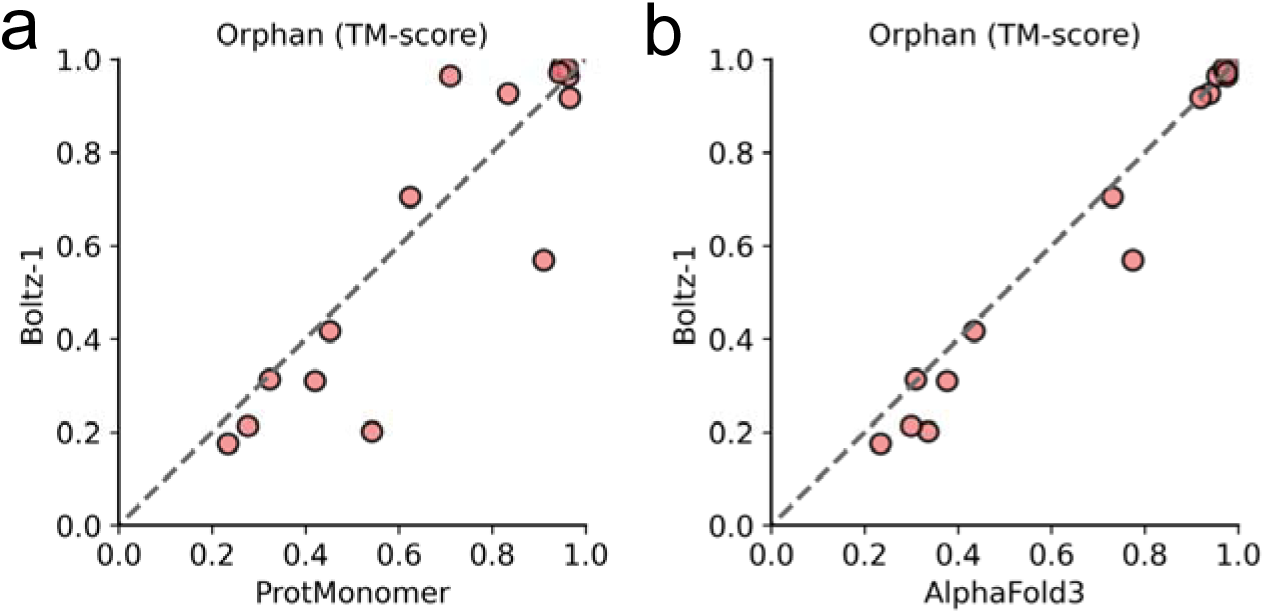
Pairwise comparisons of structures predicted by Boltz-1 (y-axis) with ProtMonomer. (**a,** x-axis) and AlphaFold3 (**b,** x-axis) on Orphan proteins, evaluated using TM-score.

**Figure S4:**
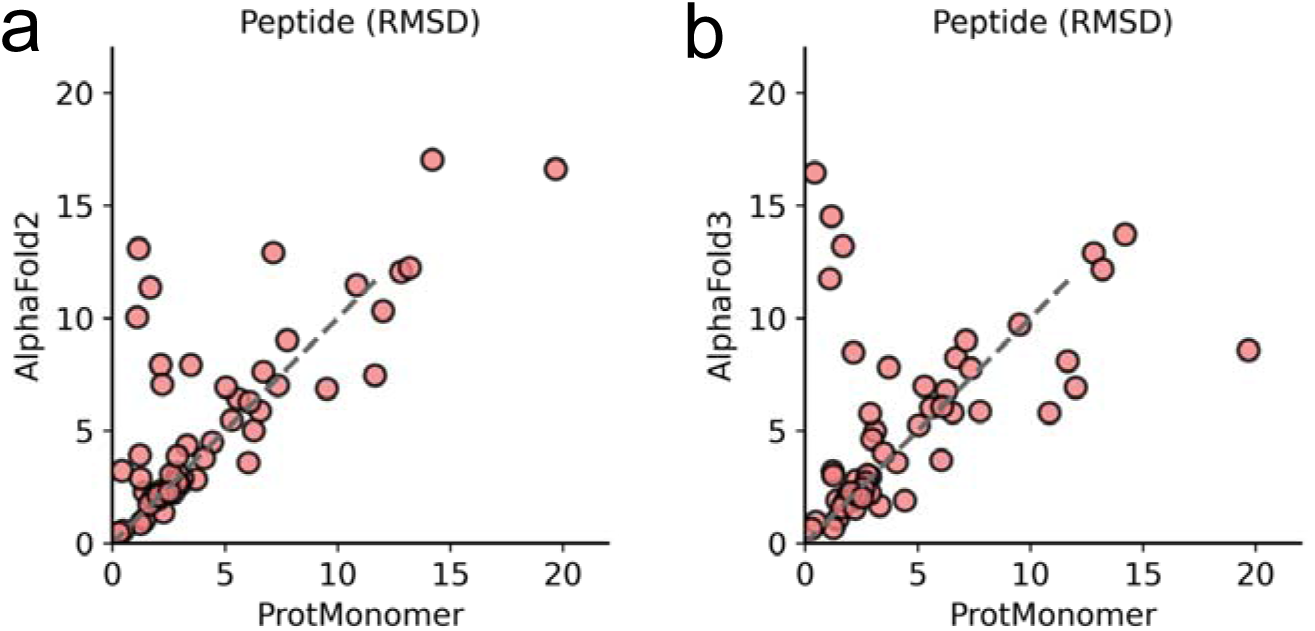
Pairwise comparison of ProtMonomer (x-axis) predictions with AlphaFold2. (**a,** y-axis) and AlphaFold3 (**b,** y-axis) on the peptide dataset, evaluated by RMSD.

**Figure S5:**
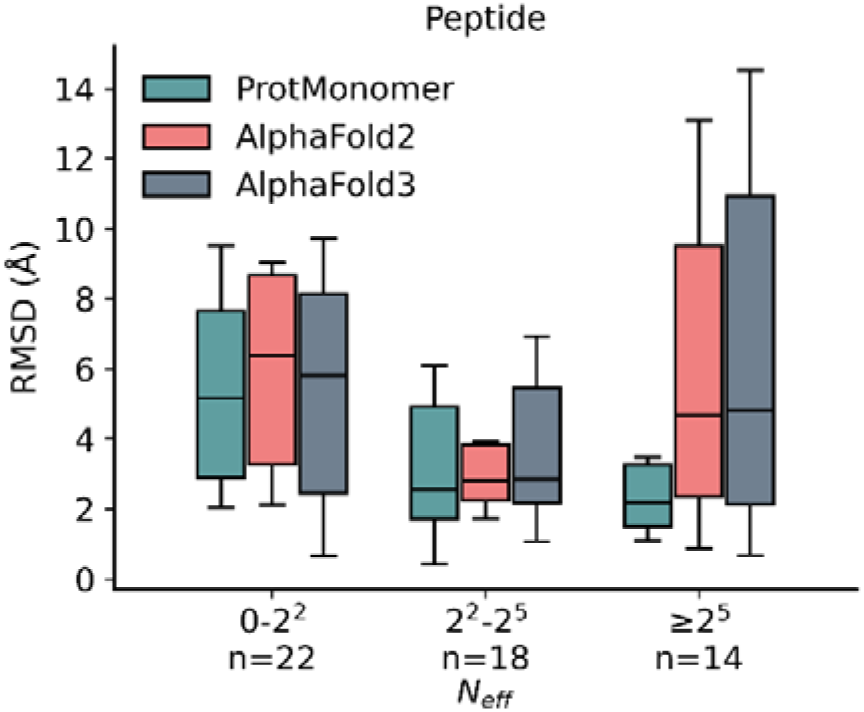
Performance of ProtMonomer, AlphaFold2 and AlphaFold3 as a function of on the Peptide dataset. Boxes indicate the interquartile range (25th–75th percentiles), center lines indicate the median and *n* denotes the number of proteins in each interval.

**Figure S6:**
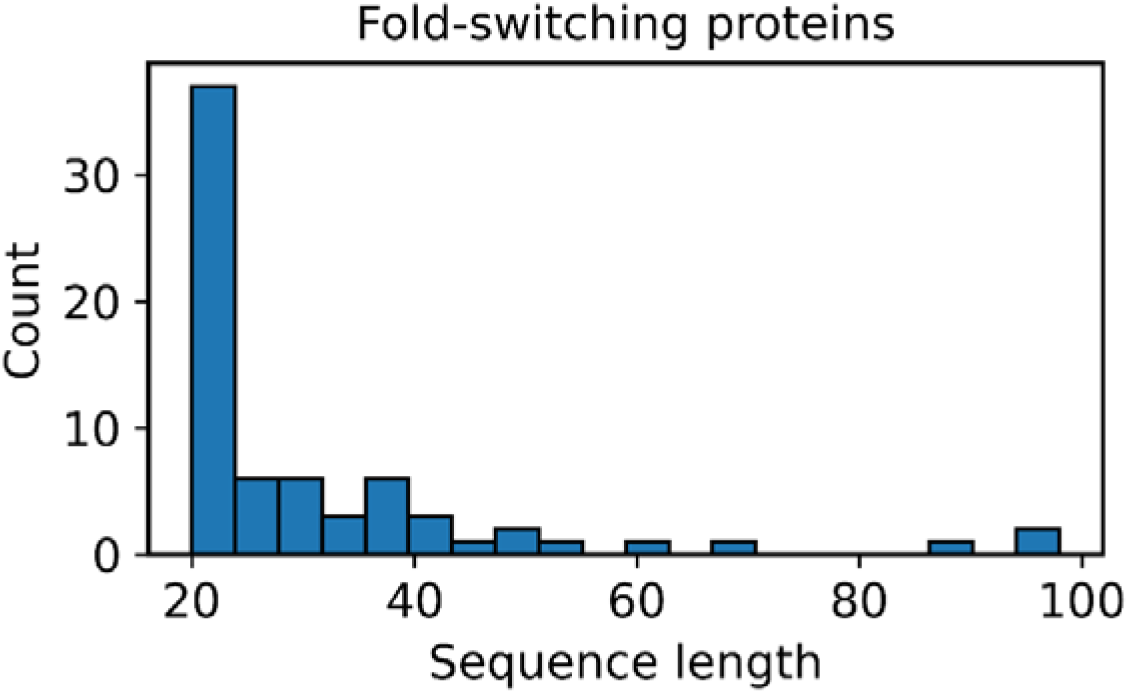
Length distribution of fold-switching proteins.

**Figure S7:**
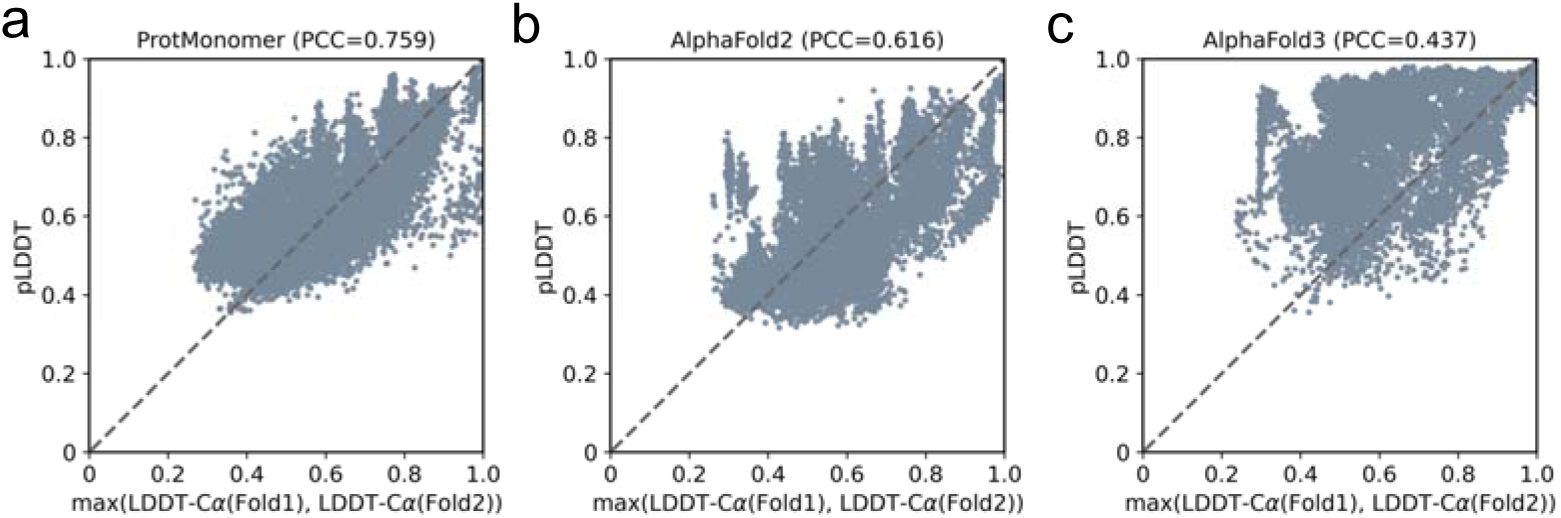
Pairwise comparison of pLDDT (y-axis) with LDDT-Cα (x-axis) for structures predicted by ProtMonomer. (**a**), AlphaFold2 (**b**) and AlphaFold3 (**c**) on the fold-switching dataset. LDDT-Cα(Fold1) denotes the LDDT-Cα value of predicted structures compared to Fold1. LDDT-Cα(Fold2) denotes the LDDT-Cα value of predicted structures compared to Fold2.

**Figure S8:**
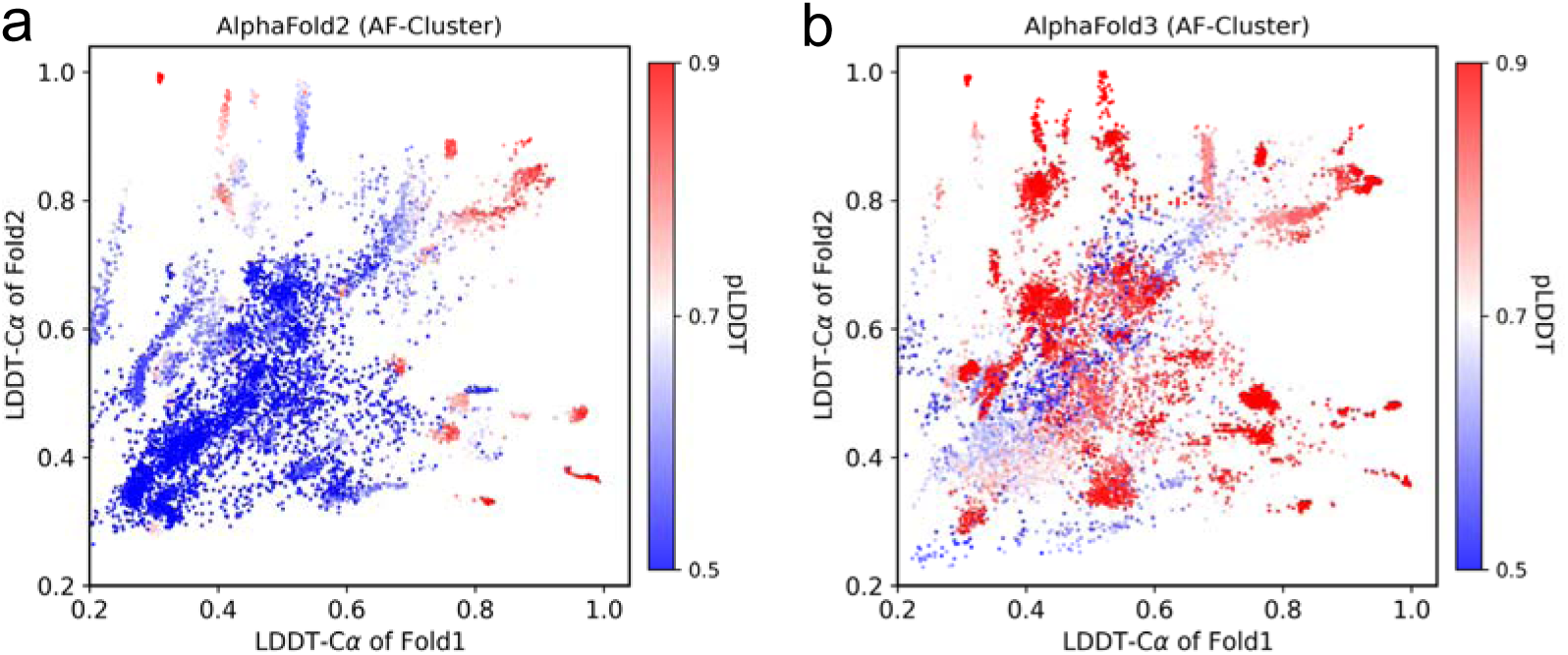
Pairwise comparison of true LDDT-Cα values relative to Fold1 (x-axis) and Fold2 (y-axis) experimental conformations, based on structures predicted by AlphaFold2. (**a**) and AlphaFold3 (**b)**, colored by pLDDT.

**Figure S9:**
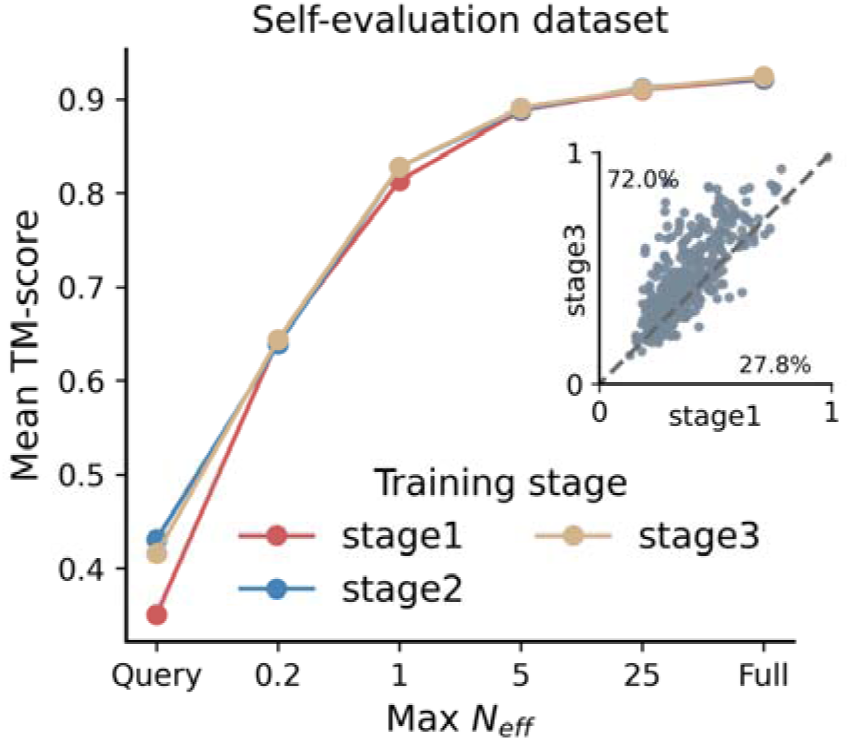
Mean TM-scores of models obtained at different training stages on the self-evaluation dataset across varying inference MSA depths. The inset shows pairwise comparisons of TM-scores for predictions from the stage 1 model (x-axis) and the stage 3 model (y-axis) on the self-evaluation dataset using only the query sequence as input.

**Figure S10:**
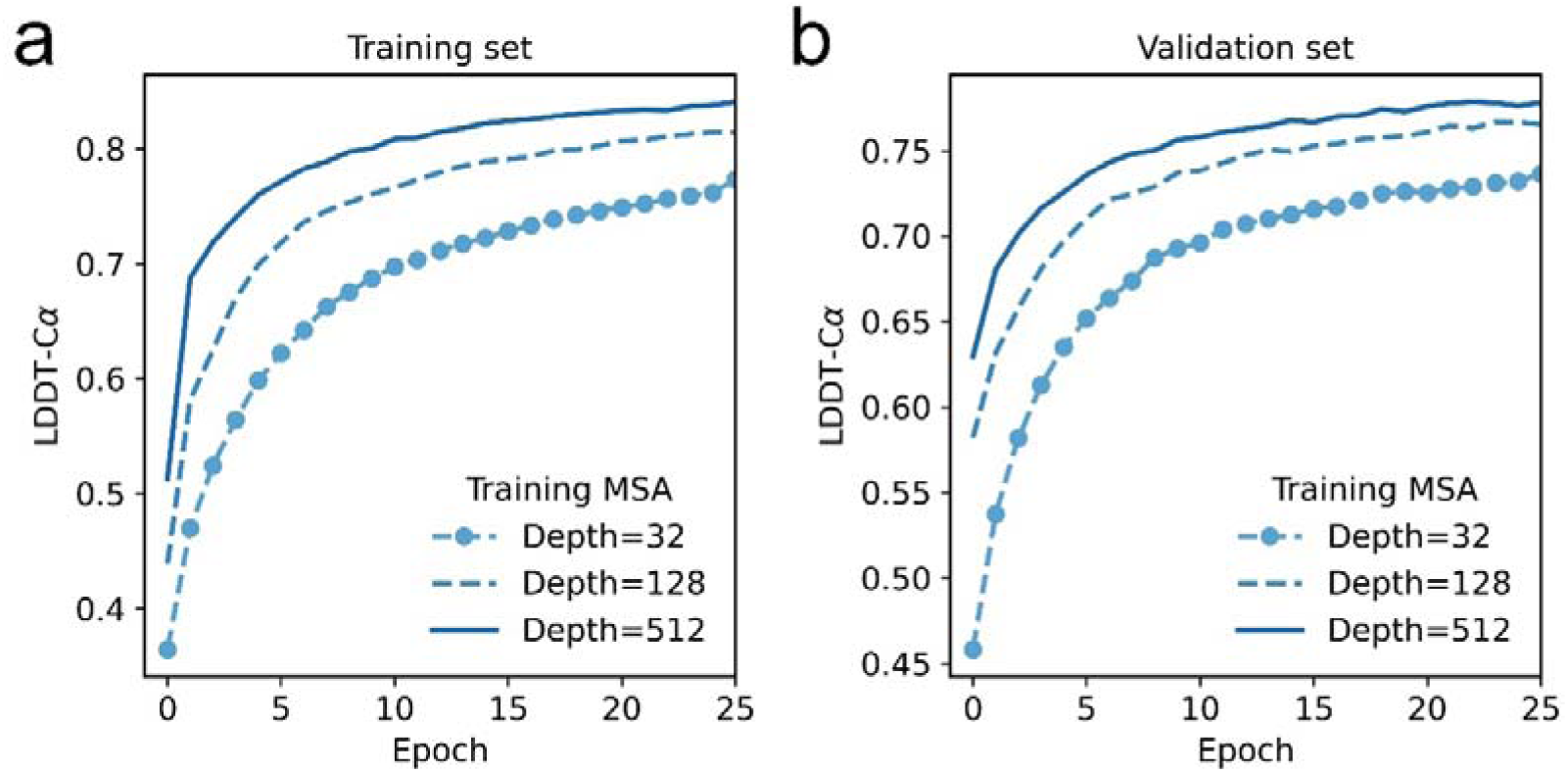
Training curves of models trained with MSA depths of 32, 128 and 512 (a: results on the training set, b: results on the validation set).

### Tables

**Table S1.** Percentage of structures predicted by ProtMonomer, AlphaFold2, and AlphaFold3 falling within different TM-score ranges for the 45 CASP15 targets.

| Method | TM-score bins |  |  |
| --- | --- | --- | --- |
|  | 0-0.5 | 0.5-0.8 | 0.8-1 |
| ProtMonomer | 8.8% | 15.5% | 75.6% |
| AlphaFold2 | 15.6% | 15.6% | 68.8% |
| AlphaFold3 | 13.3% | 17.7% | 68.8% |

**Table S2.** Mean TM-scores of ProtMonomer, AlphaFold2 and AlphaFold3 on the 17 challenging CASP15 targets. For the five-prediction setting, the prediction with the highest pLDDT score for each protein was selected for evaluation.

| Number of predictions | Method | pLDDT-selected |
| --- | --- | --- |
| 1 | ProtMonomer | 0.652 |
|  | AlphaFold2 | 0.585 |
|  | AlphaFold3 | 0.603 |
| 5 | ProtMonomer | 0.659 |
|  | AlphaFold2 | 0.608 |
|  | AlphaFold3 | 0.601 |

**Table S3.** Percentage of structures predicted by ProtMonomer, AlphaFold2, and AlphaFold3 falling within different TM-score ranges for the 866 recently released experimental structures.

| Method | TM-score bins |  |  |
| --- | --- | --- | --- |
|  | 0-0.5 | 0.5-0.8 | 0.8-1 |
| ProtMonomer | 2.7% | 12.0% | 85.2% |
| AlphaFold2 | 4.6% | 11.2% | 84.2% |
| AlphaFold3 | 5.2% | 11.6% | 83.1% |

**Table S4.** Percentage of structures predicted by ProtMonomer, AlphaFold2, and AlphaFold3 falling within different TM-score ranges for the 176 Recent PDB targets.

| Method | TM-score bins |  |  |
| --- | --- | --- | --- |
|  | 0-0.5 | 0.5-0.8 | 0.8-1 |
| ProtMonomer | 13.6% | 59.1% | 27.2% |
| AlphaFold2 | 22.7% | 55.1% | 22.2% |
| AlphaFold3 | 25.6% | 57.3% | 17.0% |

**Table S5.** Median TM-scores of structures predicted by ProtMonomer, AlphaFold2, and AlphaFold3 for Recent PDB targets grouped by different *N_eff_* intervals.

| Method | $N_{eff}$ interval | | | | | |
| --- | --- | --- | --- | --- | --- | --- |
| | $0 - 2^1$ | $2^1 - 2^3$ | $2^3 - 2^5$ | $2^5 - 2^7$ | $2^7 - 2^9$ | $\geq 2^9$ |
| ProtMonomer | 0.594 | 0.652 | 0.751 | 0.719 | 0.764 | 0.780 |
| AlphaFold2 | 0.451 | 0.610 | 0.687 | 0.699 | 0.699 | 0.771 |
| AlphaFold3 | 0.465 | 0.555 | 0.643 | 0.643 | 0.677 | 0.727 |

**Table S6.** Median RMSD (Å) of structures predicted by ProtMonomer, AlphaFold2, and AlphaFold3 for Peptide targets grouped by different *N_eff_* intervals.

| Method | $N_{eff}$ interval | | |
| --- | --- | --- | --- |
| | $0 - 2^2$ | $2^2 - 2^5$ | $\geq 2^5$ |
| ProtMonomer | 5.17 | 2.55 | 2.2 |
| AlphaFold2 | 6.37 | 2.82 | 4.69 |
| AlphaFold3 | 5.81 | 2.86 | 4.81 |

**Table S7.** Success rates of ProtMonomer, AlphaFold2, and AlphaFold3 using the AF-Cluster sampling strategy on fold-switching proteins at different RMSD thresholds between the two experimental conformations.

| RMSD<br>between two<br>states | Method |  |  |
| --- | --- | --- | --- |
|  | ProtMonomer | AlphaFold2 | AlphaFold3 |
| >2 Å | 54.5% | 13.6% | 12.1% |
| >3 Å | 54.7% | 14.1% | 12.5% |
| >4 Å | 54.7% | 14.1% | 12.5% |
| >5 Å | 50.8% | 11.8% | 8.47% |
| >6 Å | 43.7% | 8.33% | 8.33% |
| >7 Å | 31.5% | 7.89% | 5.26% |
| >8 Å | 29.0% | 6.45% | 6.45% |
| >9 Å | 25.9% | 7.41% | 7.41% |
| >10 Å | 22.7% | 4.55% | 9.09% |
| >11 Å | 13.3% | 0% | 0% |

**Table S8.** Training stages and hyperparameters of ProtMonomer.

|  | Stage 1 | Stage 2 | Stage 3 |
| --- | --- | --- | --- |
| MSA interval | [(1, 32),<br>(32, 128),<br>(128, 512),<br>(512, 2048)] | [(1,8), (8,64),<br>(64,512),<br>(512,2048), (2048,<br>16384)] | [(1,4), (4,16),<br>(16,64), (64,256),<br>(256, 1024),<br>(1024, 4096),<br>(4096, 16384)] |
| Sampling probability | [0.25, 0.25, 0.25,<br>0.25] | [0.2, 0.2, 0.2, 0.2,<br>0.2] | [0.3, 0.2, 0.1, 0.1,<br>0.1, 0.1, 0.1] |
| Warm-up steps | 1000 | 0 | 0 |
| Max residues input | 256×1280 | 340×8192 | 340×8192 |
| Learning rate | 0.001 | 0.0005 | 0.0001 |
| Epochs | 100 | 20 | 30 |
| Distillation probability | 0.75 | 0.5 | 0 |
| Crop length | 256 | 768 | 768 |
| Proteins per epoch | 50000 | 50000 | 32009 |

## Model architecture

### Main framework

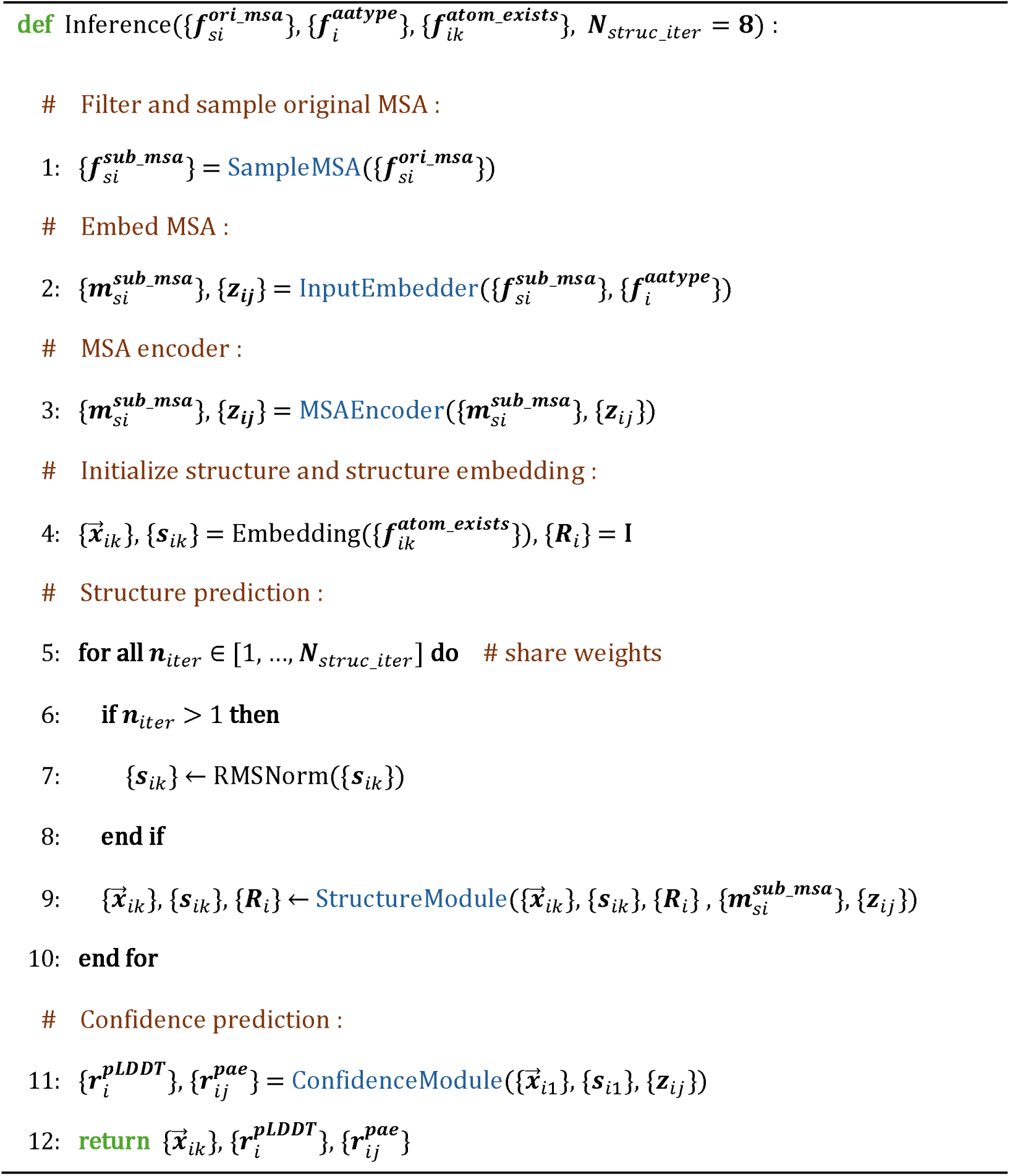
Algorithm 1 Inference.

ProtMonomer takes an MSA as input and outputs predicted all-atom coordinates, per-residue pLDDT scores, and predicted aligned error (pAE) values. The input MSA is processed sequentially by five modules. First, SampleMSA selects a high-quality sub-MSA from the original alignment, substantially reducing computational cost while preserving predictive accuracy. InputEmbedder then encodes the residue-relative positional information and embeds the sampled MSA. Next, MSAEncoder extracts MSA representations and infers inter-residue interactions from the embedded MSA. Based on these representations, StructureModule iteratively predicts and refines the atomic coordinates. In addition to the MSA-derived representations, StructureModule maintains the atomic coordinates 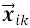, representing the coordinates of atom k in residue i, and an orientation matrix *R_i_*, which defines the local coordinate frame of residue i. These atomic coordinates are randomly initialized, whereas the orientation matrices are initialized as identity matrices. Finally, the ConfidenceModule integrates the structure and structure representations from the StructureModule with the pair representations generated by the MSAEncoder to estimate per-residue pLDDT scores and predicted aligned error (pAE) values. Here, k=1 denotes the Cα atom.

### SampleMSA (Inference)

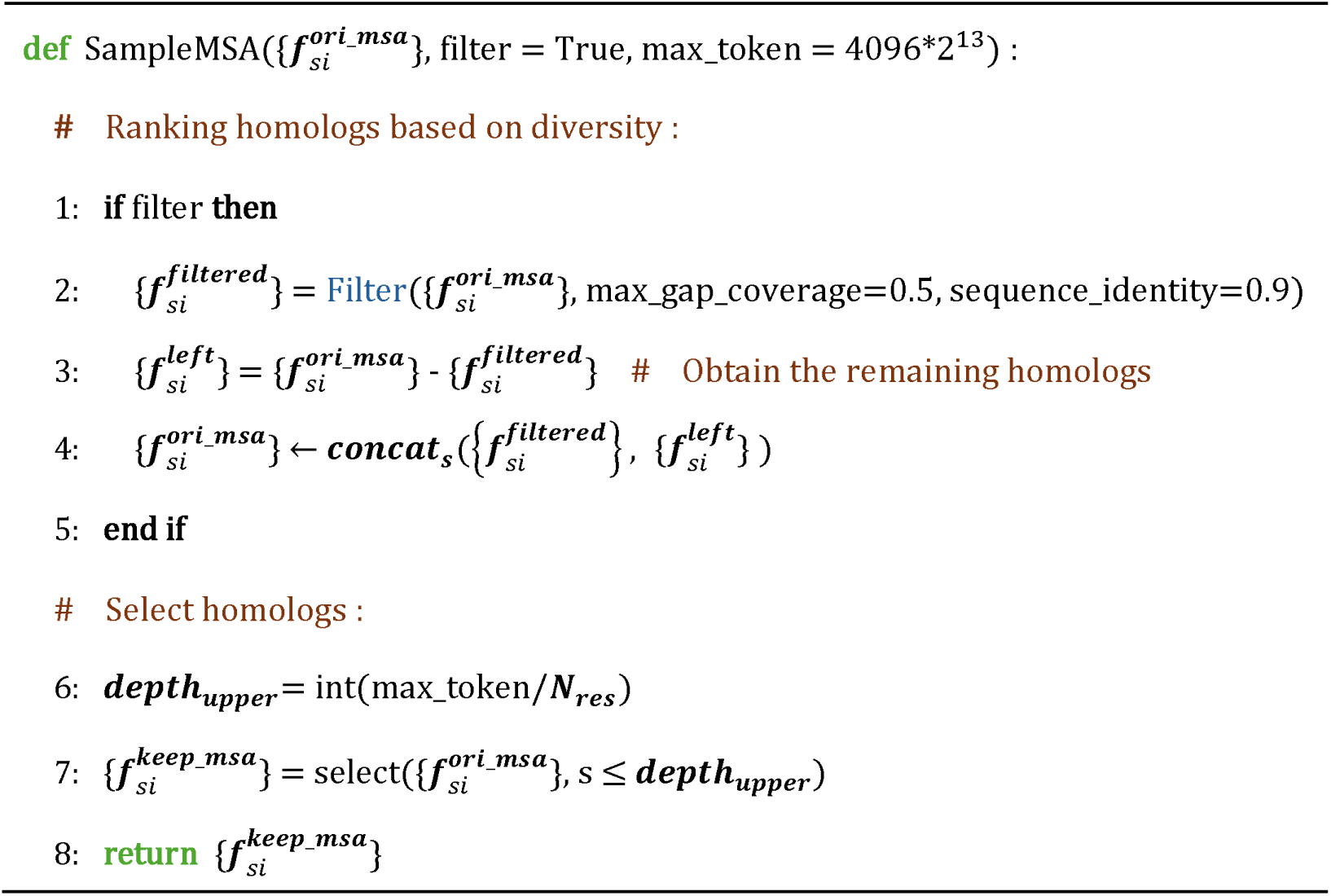
Algorithm 2 SampleMSA (Inference)

The “Filter” function is implemented using HHFilter. When enabled, it prioritizes a maximally diverse subset of homologous sequences to construct the sub-MSA. The max_token parameter controls the maximum number of residues retained in the output alignment. For a protein of 512 residues, ProtMonomer can successfully generate atomic coordinates using up to 65,536 homologous sequences on an NVIDIA H100 PCIe GPU. The variable 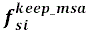 denotes amino acid type at residue i in the s-th homologous sequence retained in the filtered MSA.

### SampleMSA (Training)

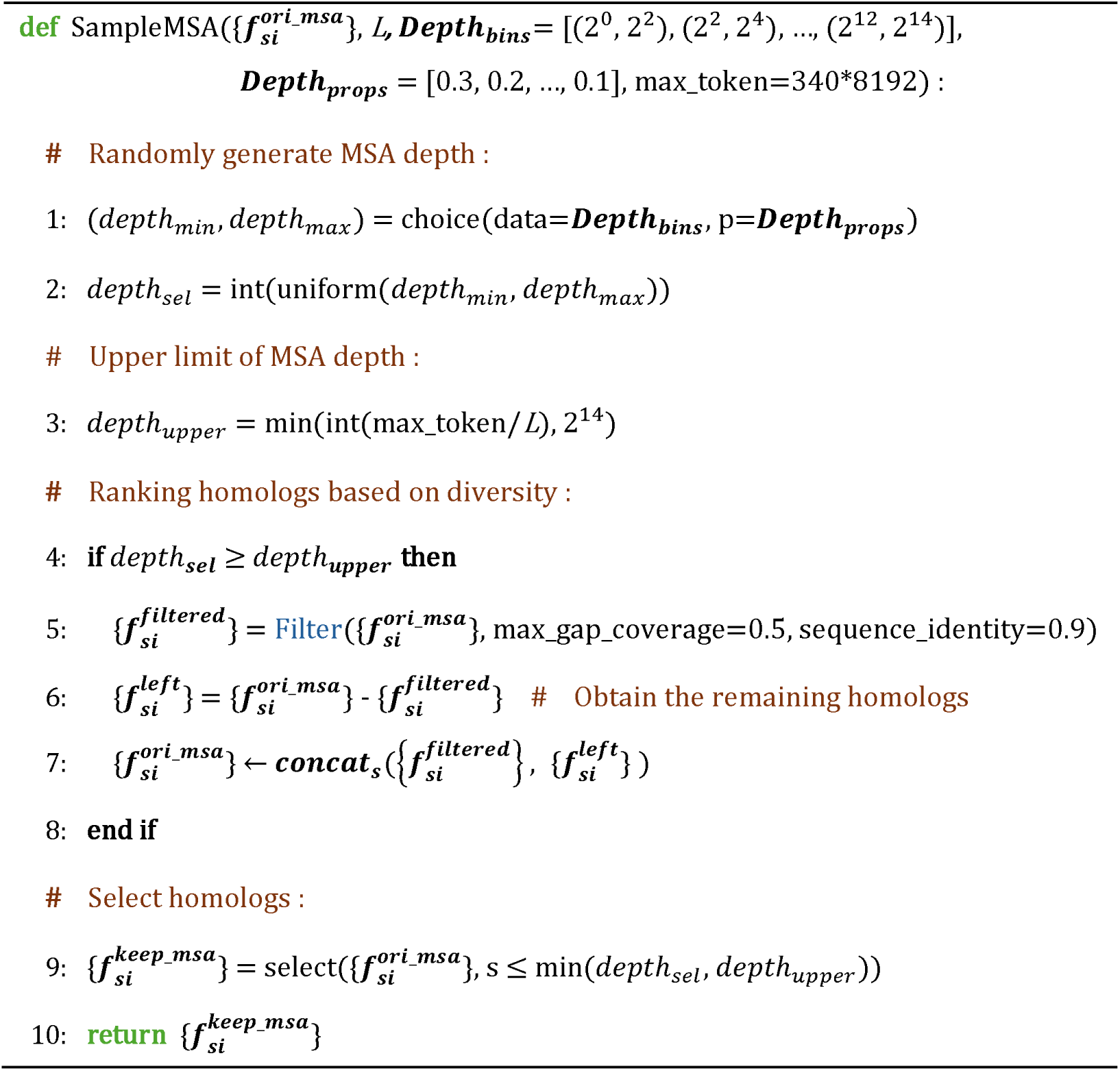
Algorithm 3 SampleMSA (Training)

*Depth_bins_* denotes the intervals spanning different MSA depths, L denotes the sequence length, and *Depth_props_* denotes the sampling probability assigned to each interval.

### InputEmbedder

The InputEmbedder takes the query sequence and sub-MSA as inputs and produces two types of embeddings. The sub-MSA embedding {*m_si_*} is obtained by applying an embedding layer to the sub-MSA and has shape (N, L, 256), where N is the number of homologous sequences and L is the sequence length. The pair embedding {*z_i,j_*} is constructed by combining the embedded query sequence (from an embedding layer) with an embedded relative positional encoding of residue pairs following the approach used in AlphaFold2, and has shape (L, L, 128).

### MSAEncoder

The MSAEncoder consists of 32 layers of a modified Evoformer architecture, in which LayerNorm is replaced with RMSNorm to slightly reduce computational cost.

### StructureModule

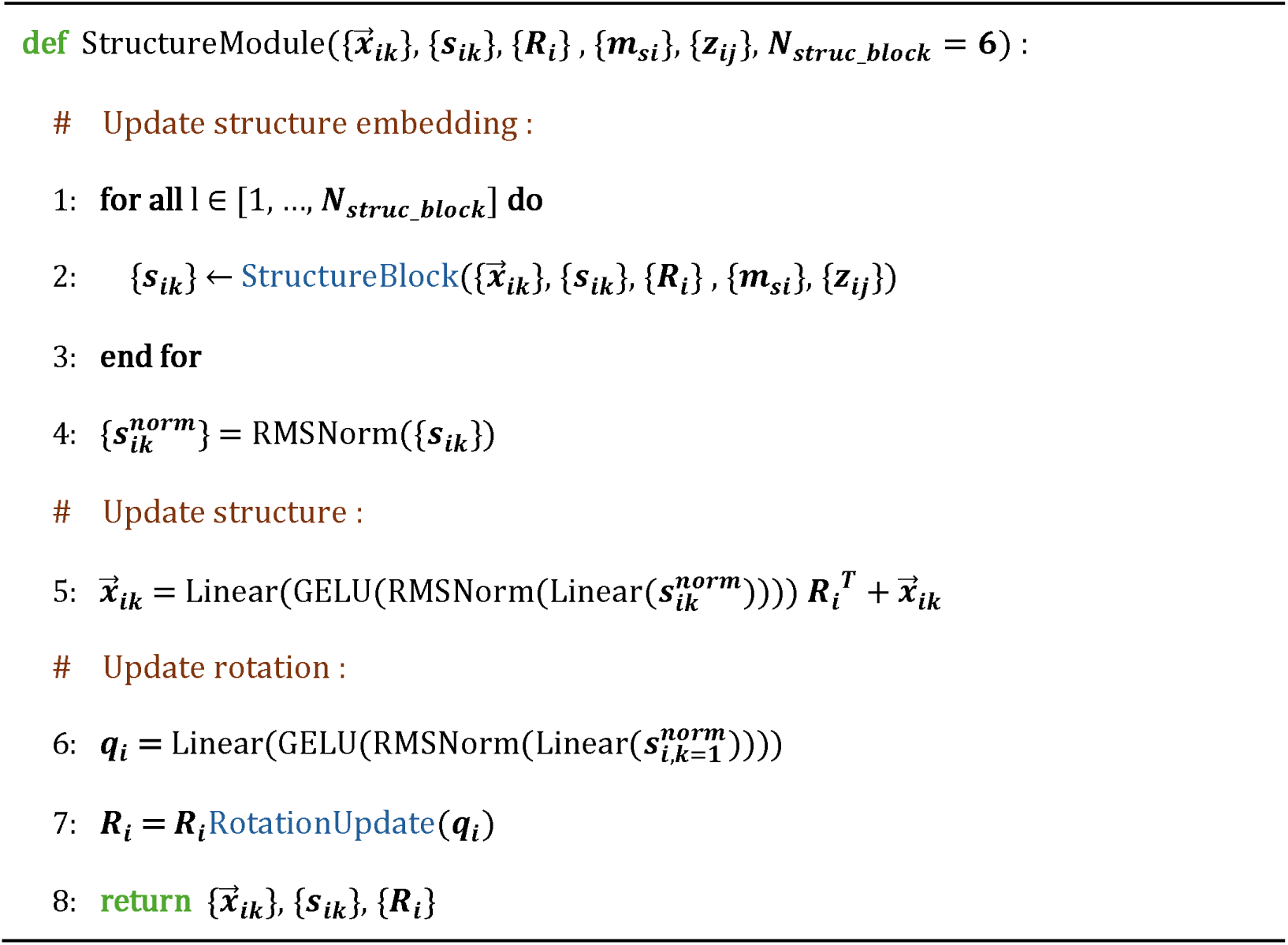
Algorithm 4 StructureModule.

The StructureModule, consisting of six structure blocks, predicts and iteratively refines the all-atom protein structure based on the MSA and pair representations. Here, (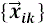 denotes atom coordinates, with a shape of (L, 37, 3). L denotes sequence length and 37 corresponds to the set of atom types used to represent standard amino acids. {*s_ik_*} denotes the structure representation, with a shape of (L, 37, 256). {*R_i_*} denotes the local coordinate frame defined on each residue, with a shape of (L, 3, 3). {*z_i,j_*} denotes the pair representation, with a shape of (L, L, 128).

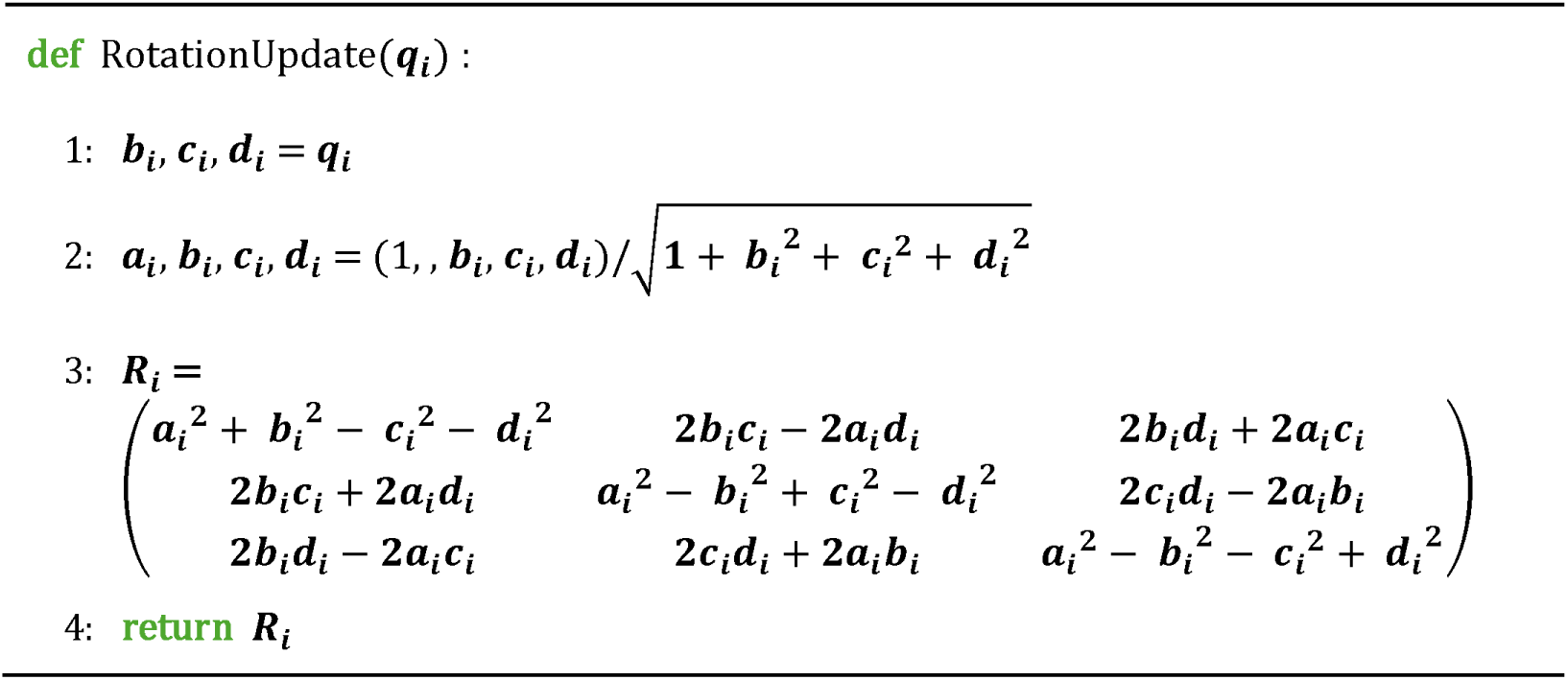
Algorithm 5 Rotation update.

### StructureBlock

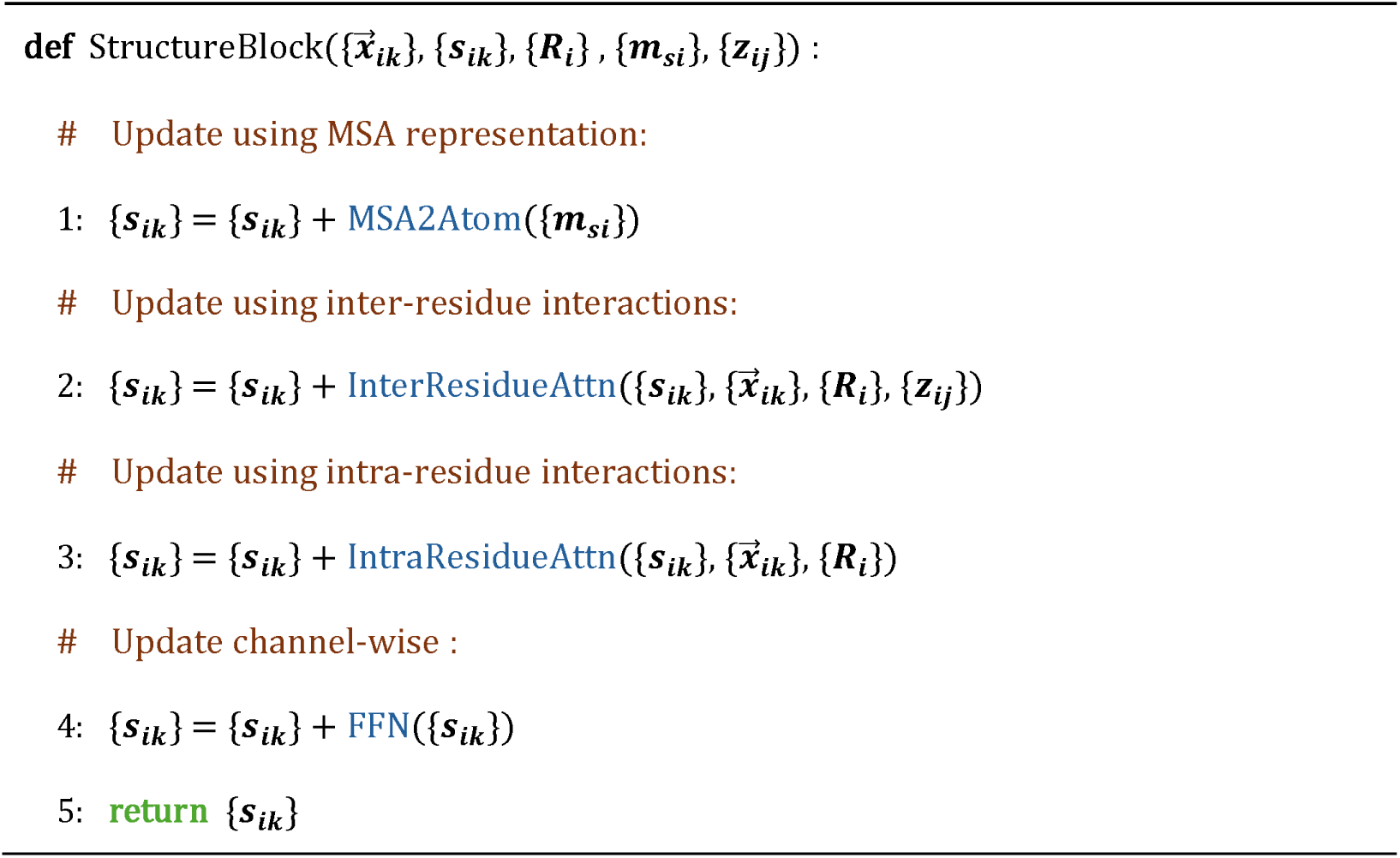
Algorithm 6 StructureBlock.

Each structure block receives the MSA representation and pair representation from the MSA encoder, the predicted structure from the previous iteration and the current structure representation as inputs.

### MSA2Atom

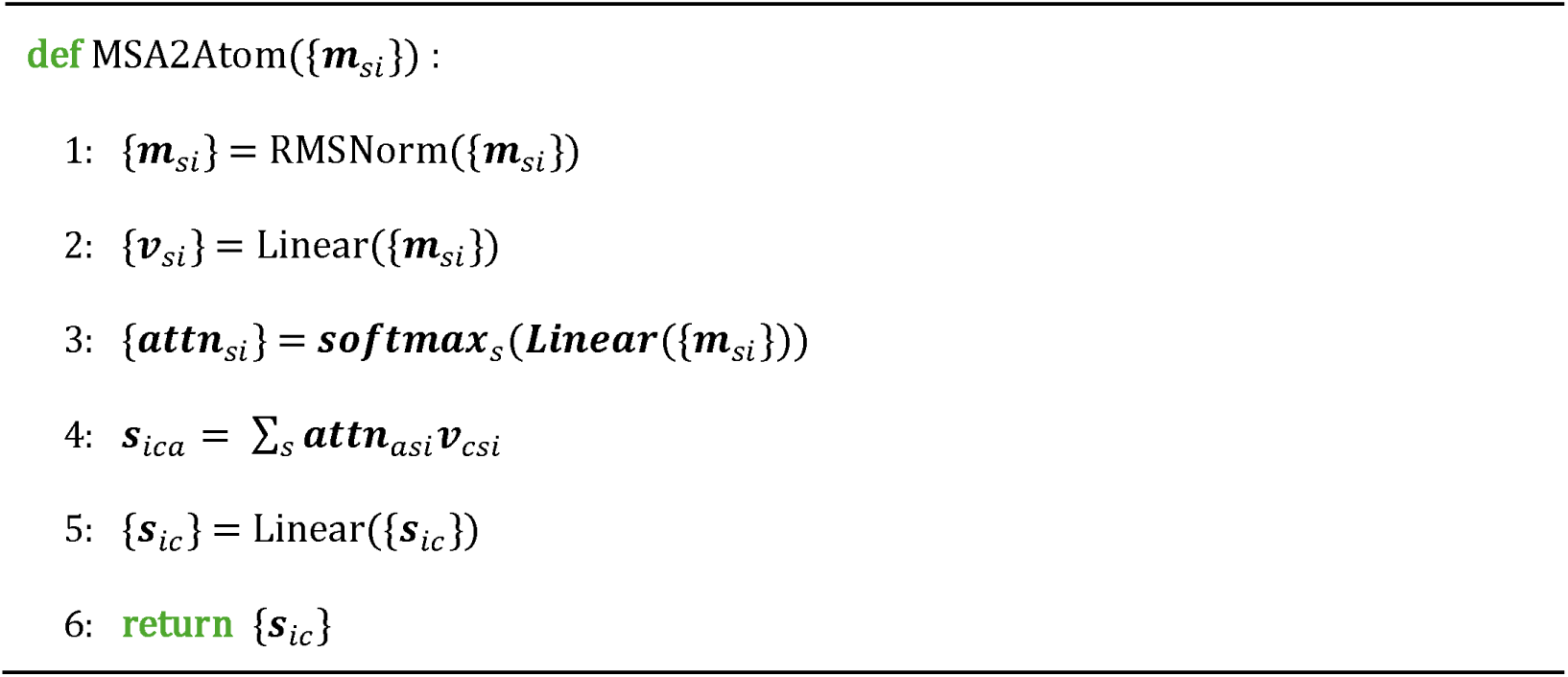
Algorithm 7 MSA2Atom.

MSA2Atom transforms the MSA representation into an atom-level structure representation. {*v_si_*} denotes an intermediate representation derived from the MSA, with a shape of (N, L, 37), where N is the number of homologous sequences and L is the sequence length. *s_aic_* denotes the resulting atom-level representation, with shape (L, 37, 256).

### InterResidueAttn

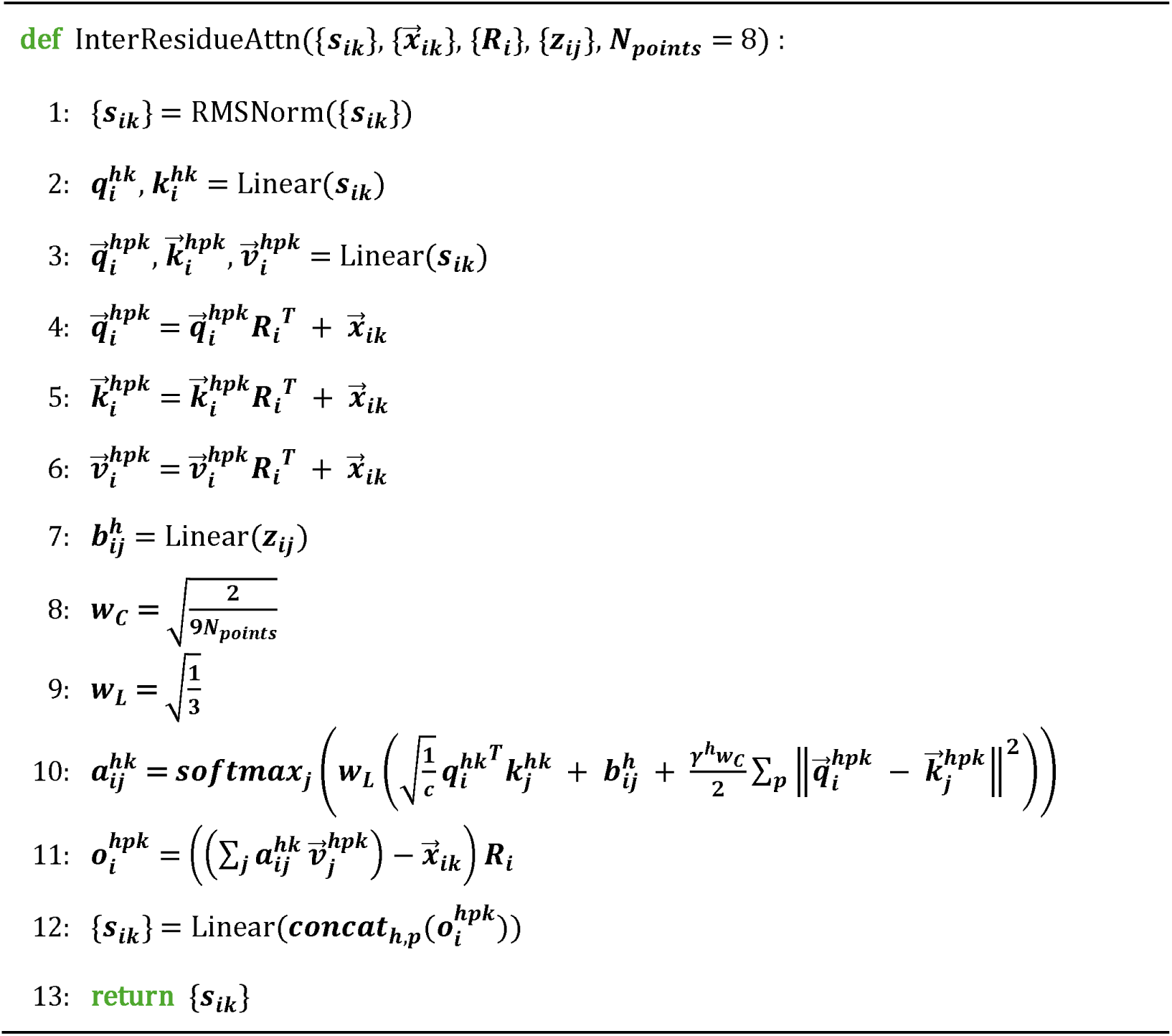
Algorithm 8 InterResidueAttn.

InterResidueAttn is used to update the structure representation with inter-residue interactions. It is based on the invariant point attention (IPA) module introduced in AlphaFold2 but is substantially simplified and computes residue–residue interactions independently for each atom channel.

### IntraResidueAttn

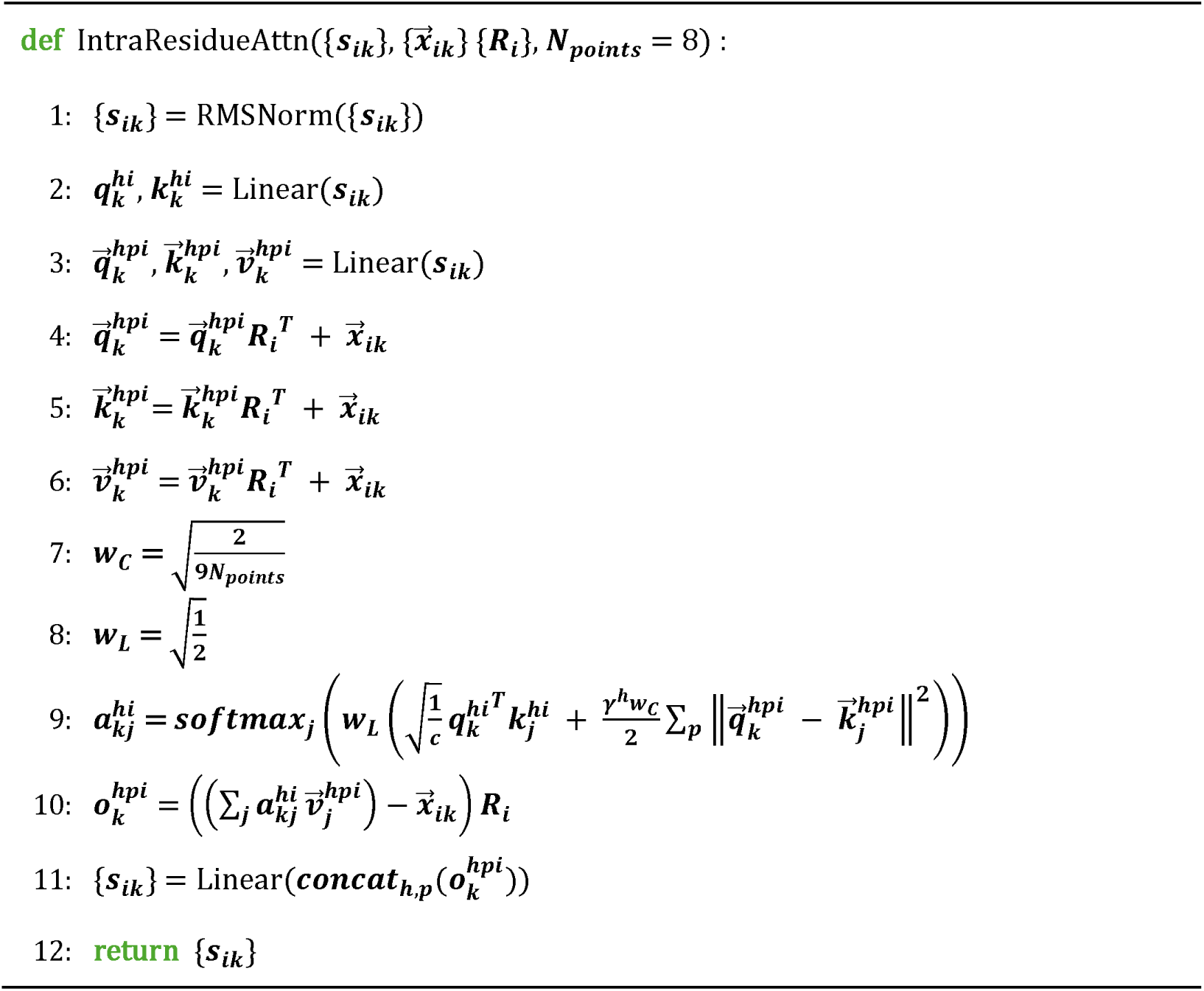
Algorithm 9 IntraResidueAttn.

IntraResidueAttn is designed to update the structure representation by modeling intra-residue interactions. Compared with InterResidueAttn, it does not require the pair representation as an input.

### FeedForwardNetwork

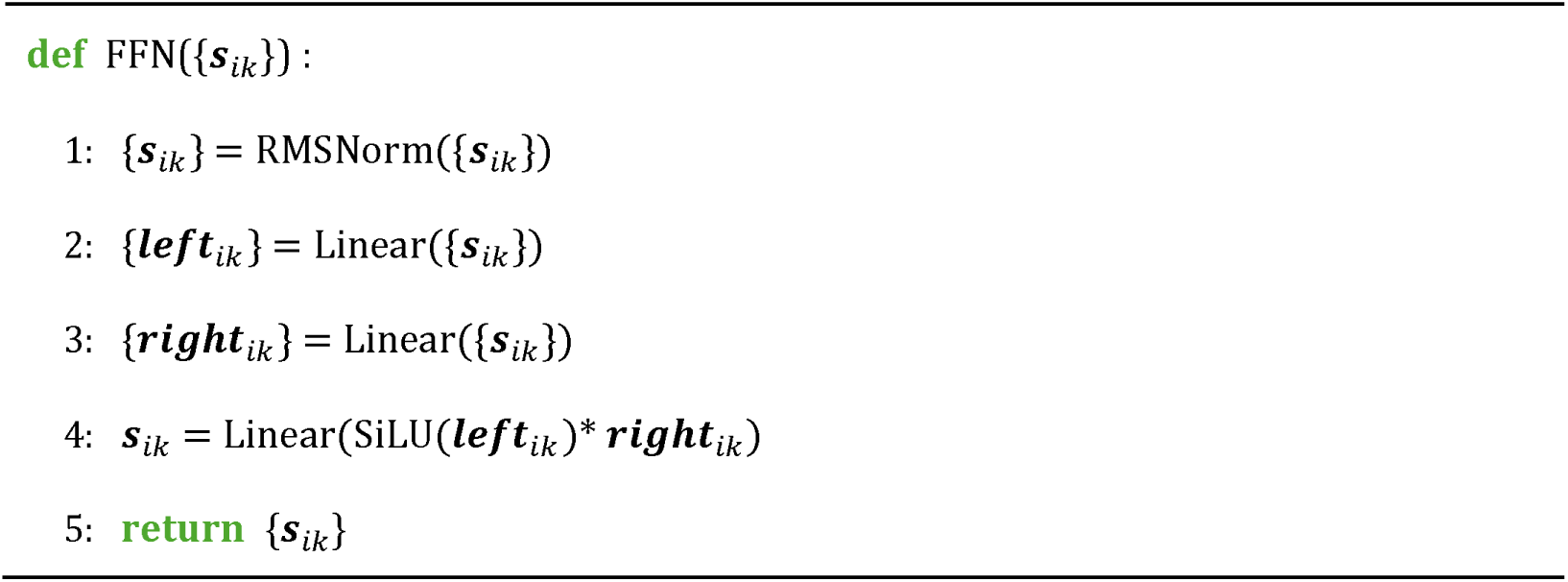
Algorithm 10 FeedForwardNetwork.

The FFN module applies a channel-wise transformation to further refine the structure representation.

### MaskedMSAHead

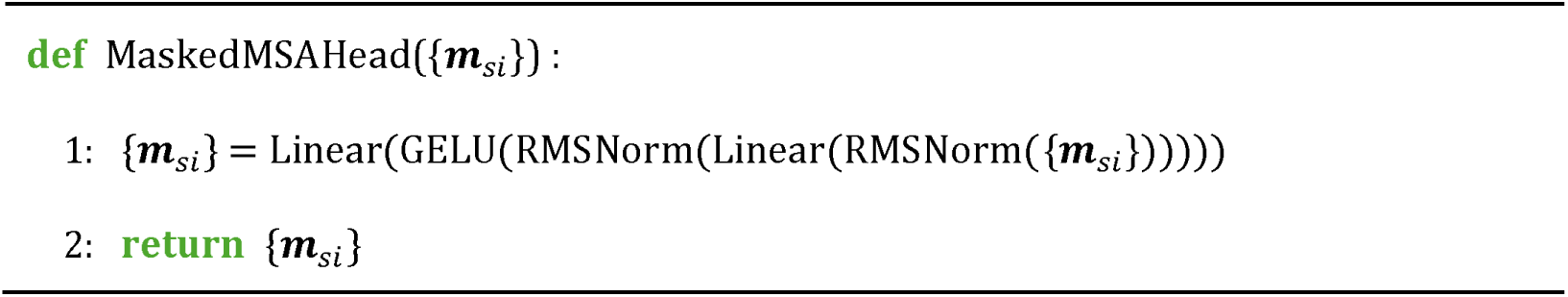
Algorithm 11 Masked MSA head.

MaskedMSAHead predicts the identities of masked residues in the input MSA using the MSA representations generated by the MSAEncoder.

### DistogramHead

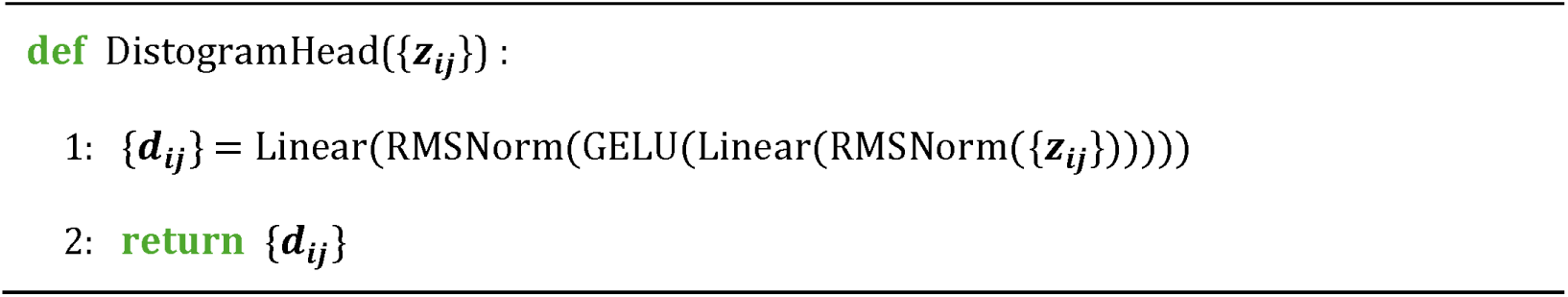
Algorithm 12 Distogram Head.

DistogramHead predicts pairwise Cβ–Cβ distances from the pair representations generated by the MSAEncoder. The distances are discretized into 64 bins spanning 2.3125–21.6875 Å^15^.

### ConfidenceHead

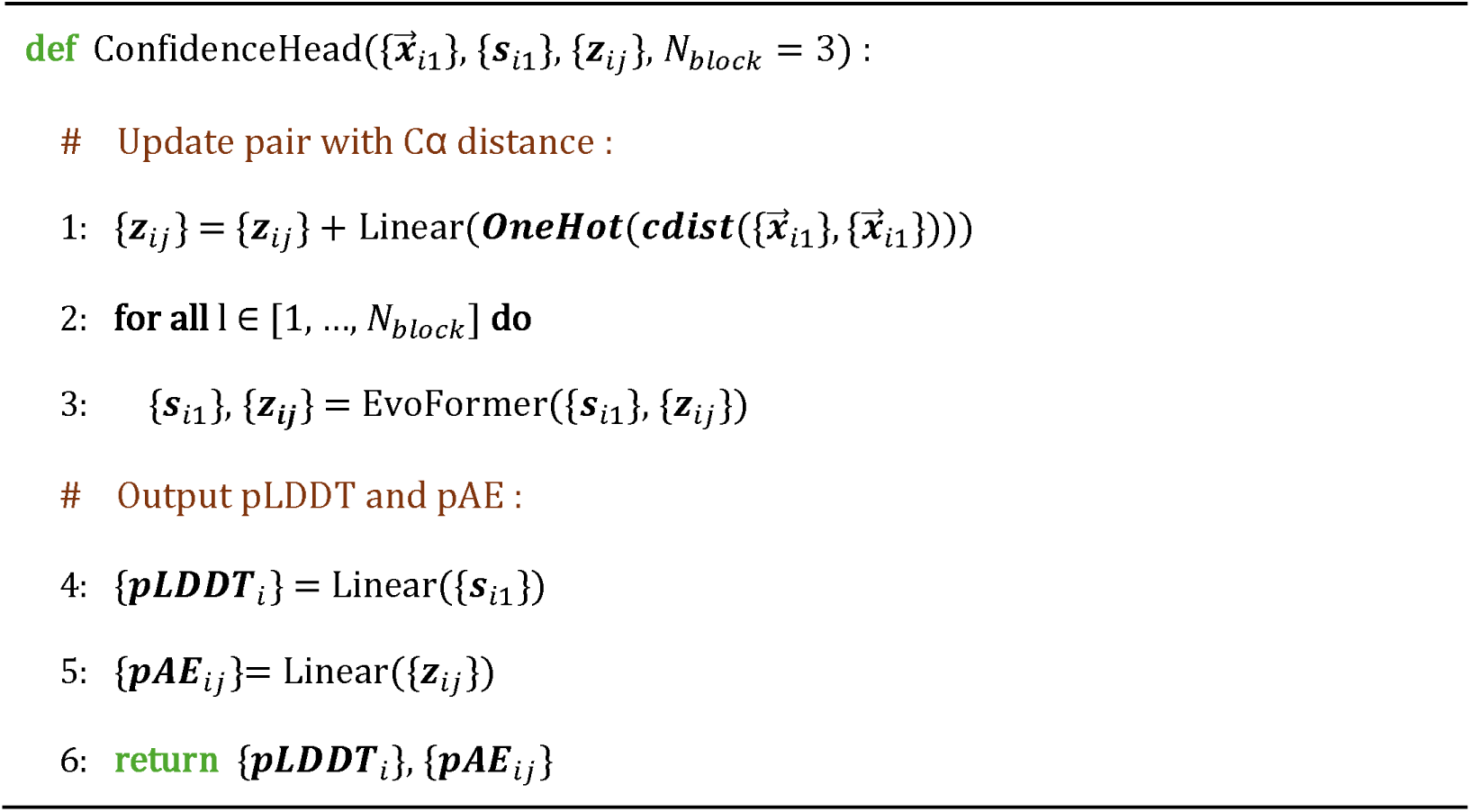
Algorithm 13 ConfidenceHead.

ConfidenceHead estimates per-residue pLDDT scores and predicted aligned error (pAE) values for the structures produced by the StructureModule. The cdist function is used to compute the pairwise Cα–Cα distance matrix, and the resulting distances are discretized into 16 bins spanning 3.25–20.75 Å and encoded using one-hot representations. The pLDDT output has shape (L, 50), corresponding to 50 discrete confidence bins for each residue, whereas the pAE output has shape (L, L, 64), corresponding to 64 discrete error bins for each residue pair.

### All-atom and all-frame FAPE Loss

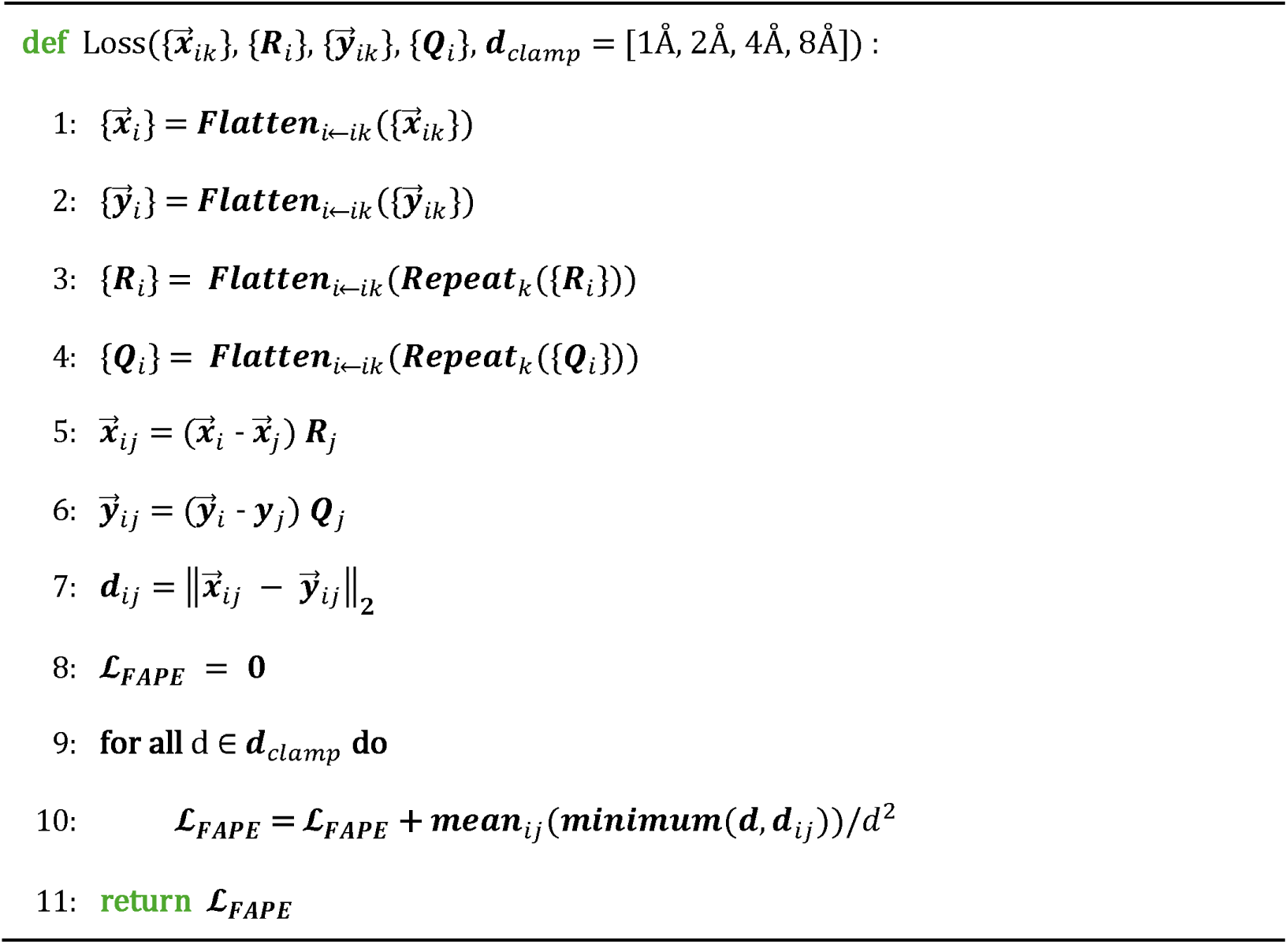
Algorithm 14 all-atom and all-frame FAPE Loss.

The coordinates of the predicted structure 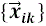 and the experimental structure 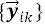 both have shape (L, 37, 3). The predicted local coordinate system {*R_i_*} has shape (L, 3, 3). The local coordinate system of the experimental structure, {*Q_i_*}, also has shape (L, 3, 3) and is constructed for each residue from the backbone N, Cα, and C atoms using Gram–Schmidt orthogonalization, following the procedure used in AlphaFold2.

